# Changes amid constancy: flower and leaf microbiomes along land use gradients and between bioregions

**DOI:** 10.1101/2020.03.31.017996

**Authors:** Paul Gaube, Robert R. Junker, Alexander Keller

**Affiliations:** Department of Bioinformatics, University of Würzburg, D-97074 Würzburg, Germany; Center for Computational and Theoretical Biology, University of Würzburg, D-97074 Würzburg, Germany; Department of Biosciences, University of Salzburg, A-5020 Salzburg, Austria; Evolutionary Ecology of Plants, Faculty of Biology, Philipps-University Marburg, D-35032 Marburg, Germany

**Keywords:** *Ranunculus acris*, *Trifolium pratense*, plant-associated bacteria, phyllosphere, microbial ecology, metabarcoding

## Abstract

Microbial communities inhabiting above-ground parts of plants affect their host’s development, fitness and function. Although studies on plant-associated microbes are of growing interest, environmental drivers of flower microbiomes in particular are poorly characterized. In this study, we investigated flower and leaf epiphytic bacterial microbiomes of *Ranunculus acris* and *Trifolium pratense* using metabarcoding of 16S ribosomal DNA in three German bioregions and along land-use intensity gradients. Our data suggests that the structures of bacterial communities clearly differed between plant species and tissue types. Also, floral bacterial communities of *R. acris* showed higher variability in comparison to *T. pratense*. Bacteria usually associated with pollinators were found solely in flower samples, while such usually associated with the rhizosphere were only present in high abundances on leaves. We identified Pseudomonadaceae, Enterobacteriaceae and Sphingomonadaceae as the most abundant taxa on flowers, while Sphingomonadaceae, Methylobacteriaceae and Cytophagaceae dominated bacterial communities on leaves. We found strong bacterial turnover already for short geographic distances, which however did not increase with the long distances between bioregions. High land use intensity caused phylogenetically less diverse and more homogenous bacterial communities. This was associated with a loss of rare bacterial families. Intensification of mowing and fertilization affected almost all plant associated bacterial communities, while grazing had only minor effects on bacterial structures of *T. pratense* flowers. However, dominant taxa were mostly resilient to mowing, grazing and fertilization. Despite that, we identified indicator taxa for regularly disturbed environments in flower microbiomes.

## Introduction

Above-ground plant microbiota (phyllosphere) are known for their importance for host growth, health and fitness, as particularly well studied in associations of leaves (phylloplane) (Bulgarelli et al., 2013; Compant et al., 2019; Vorholt, 2012). In recent years, it has become increasingly recognized that bacteria associated with flowers (anthosphere) are also directly related to plant health and fitness. However, there is still little known about the determinants and the composition of floral microbiomes (Aleklett et al., 2014). In comparison to leaves, flower microbiomes are less diverse and differ in their composition (Junker & Keller, 2015; Junker et al., 2011; Krimm et al., 2005; Leff et al., 2015). The most abundant bacterial phylum associated with flowers is represented by Proteobacteria, especially by Pseudomonas, Enterobacteriaceae, Sphingomonas and Methylobacterium. Further less abundant taxa belong to Actinobacteria, Bacteroidetes and Firmicutes (Aleklett et al., 2014; Junker & Keller, 2015; Rebolleda Gómez & Ashman, 2019; Steven et al., 2018; Wei & Ashman, 2018). The structures of floral microbiomes are potentially influenced by other bacterial habitats such as soils or leaves. Bacterial colonizers of flowers are likely transmitted through air and rain, vascular tissues or by animals especially by pollinators (Belisle et al., 2012; de Vega & Herrera, 2013; McFrederick et al., 2017; Vannette & Fukami, 2017). In contrast to leaves, the lifespan of fully developed flowers is only a few days. Hence, there is a limited time space of environmentally transmitted bacterial acquisition. Recently it has been shown that bacteria are transmitted between flowers by pollinators (McFrederick et al., 2017; Voulgari-Kokota et al., 2019a; Voulgari-Kokota et al., 2019b) and that flower associated bacteria can have an impact on the emission of volatiles that potentially attract pollinators (Helletsgruber et al., 2017; Penuelas et al., 2014). Further linkages of floral bacteria to pollination service have been reported, causing fitness advantages by defensive mutualism (D.-R. Kim et al., 2019), having impacts on chemical nectar traits (Vannette & Fukami, 2018) or for density-dependent negative effects on pollinator visits (Junker et al., 2014).

The microhabitats of the anthosphere differ in their morphological structures and chemical compositions, ranging from poor to nutrient rich environments. Especially the nectar but also stigmas, styles and pollen provide excellent conditions for microbial growth in form of easily degradable sugars, amino acids and floral waxes that could privilege fast-growing and highly competitive bacteria (Mercier & Lindow, 2000; Wilson & Lindow, 1994). Nevertheless, flowers harbour also a variety of active and passive antimicrobial defence mechanisms that select only for specific or well adapted bacteria (Gonzalez-Teuber et al., 2009; Harper et al., 2010; Huang et al., 2012; Junker & Tholl, 2013). The host plants benefit from selected microbial colonizers because it is likely that they ensure protection against phytopathogens and enhance stress tolerance of the plants (Pusey et al., 2011; Stockwell et al., 2010; Wilson & Lindow, 1993). Besides the availability of carbon or nitrogen sources and the acquisition of source communities, specific biogeographical conditions like climate, other abiotic factors such as surface structure, pH and moisture or biotic factors that contribute to the host immune responses are likely also important for the compositional determination of plant associated microbial communities (Aleklett et al., 2014; Compant et al., 2019; Rebolleda-Gómez et al., 2019). Considering the biogeographical distribution of plants and other organisms within a given habitat type, it is known that communities which are located close to each other, are more similar than communities that are geographically separated (Lomolino et al., 2010). Thus, we would expect a similar pattern for phyllosphere bacterial communities, becoming more dissimilar with increasing geographic distances between sites. Furthermore, it has to be considered that anthropogenic influences on intensities of land use management types like fertilization, grazing and mowing can have strong effects on soil properties and on plant pheno- and chemotypic characteristics, and therefore may also shape flower and leaf microbiomes (Estendorfer et al., 2017; Li et al., 2018; E. K. Morris et al., 2013; Schöps et al., 2018; Völler et al., 2017).

In this study we examined the epiphytic bacterial communities inhabiting healthy flowers and leaves of two plant species *Ranunculus acris* L. and *Trifolium pratense* L. by using 16S rRNA gene amplicon sequencing. The study was conducted within the framework of the German Biodiversity Exploratories project, covering continuous land-use gradients varying in degrees of mowing, grazing and fertilization. We investigated three regions that were geographically distinctly separated by at least 300 km and contained replicated plots with similar treatments. This allows to distinguish local treatments as well as short- and long-distance biogeographical effects. We first examined bacterial alpha- and beta-diversity as well as composition for samples with respect to plant tissue and species. We aimed to identify whether bacterial communities are consistent between regions and resilient to local land use intensity, or if changes are observable between bioregions and land use impacts on structure of microbial assemblages.

## Materials and Methods

### Study Side and Sample Collection

This study was conducted within the framework of the German Biodiversity Exploratories (www.biodiversity-exploratories.de), a large-scaled research project investigating the relationship between biodiversity, land-use intensity, and functional ecosystem processing. In 2017 flowers and leaves of *Ranunculus acris* L. and *Trifolium pratense* L. were sampled in grasslands of three geographically distinct regions that are managed by farmers. The three regions span latitudinally 800 km from north to south Germany and cover different landscape types (Fischer et al., 2010). The UNESCO Biosphere region Schwäbische Alb (Baden-Württemberg, ALB) is located in the southwest of Germany and is characterized by calcerous bedrock. The national park Hainich is also characterized by calcareous bedrock and in the middle of Germany (Thüringen, HAI). In the northeast of Germany is the UNESCO Biosphere region Schorfheide-Chorin (Brandenburg, SCH), which is defined by young glacially formed landscapes (Fischer et al., 2010). The experimental plots with the size of 50 m x 50 m within the grassland of the Biodiversity Exploratories are described by a land use intensity (LUI) index. The LUI index of grasslands in all three regions is spanned over a continuous gradient and is characterized by three management types: fertilization, grazing and mowing (Blüthgen et al., 2012). The plots ranged from unfertilized meadows and pastures to highly fertilized meadows and mown pastures. We classified the plots into low and high categories based on their land use management intensities (according to Estendorfer et al., 2017). Furthermore, depending on their degree of intensification, we classified the types of land use separately into the categories none, low, medium and high intensity management.

The flower and leaf samples of three individuals of both plant species per plot were sampled whenever both co-occurred on a plot. This resulted in samples from 21 plots of the Swabian-Alb, 13 plots of the Hainich and 9 plots of the Schorfheide-Chorin. All sampled plant individuals had at least 3 m distance between each other to avoid repeated sampling of the same individual. All samples were treated with 70% ethanol sterilized forceps and scissors. Samples of whole flowers (*Ranunculus acris*) and inflorescences (*Trifolium pratense*) were collected at similar stages of development and placed in 2 mL lysis tubes or in 15 mL falcon tubes containing bashing beads and 750 μL or 1500 μL of DNA/RNA Shield (Zymo Research, CA, USA), respectively. Leaf samples were obtained with sterile swabs. The leaf samples were placed in lysis tubes with 750 μL of DNA/RNA Shield™ (Zymo Research) and bashing beads. All samples were shaken for 1 minute by hand in order to obtain epiphytic bacteria and to stabilize the sampling solution. The samples were stored at approximately 8°C in the field and at −20°C until further processing in the lab.

### DNA Isolation and PCR Amplification

Genomic DNA was isolated using the ZymoBIOMICS™ 96 DNA Kit (Zymo Research) according to the manufacturer’s protocol with minor modifications as follows. Initially all samples were vortexed for 10 min at maximum speed and centrifuged for 5 min at 12,000x g. Isolated DNA were stored at −20°C until further molecular analysis.

Amplifications of the 16S rRNA gene V4 region were performed with a dual-indexing strategy according to Kozich et al. (2013). We applied a dual-indexing approach that allow us to multiplex the samples (Illumina, 2016). To minimize random amplifications, we processed the PCR reactions in triplicates with 0.5 μL of template DNA and 5 μL Phusion High-Fidelity PCR Master Mix with HF Buffer (Thermo Fisher Scientific, Waltham, USA) in each reaction. In order of quality control, we used negative controls of i) DNase/RNase Free Water (Zymo Research), ii) DNA/RNA Shield™ (Zymo Research) and iii) sterile swabs. A Microbial Community Standard (Zymo Research) was used as a positive control. All controls passed through the same workflow as the plant samples. Furthermore, we applied pPNA blocking primers (PNA Bio Inc., Newbury Park, USA) to the PCR reactions at a final concentration of 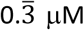 to prevent the amplification of chloroplast related sequences (Lundberg et al., 2013). PCR cycle parameters were as follows: an initial denaturation step at 95 °C for 4 min, followed by amplification steps by using 30 cycles of 95 °C for 40 s, including PNA clamping at 75 °C for 10 sec, annealing at 55 °C for 30 sec and extension at 72 °C for 60 sec. A final extension step of 72 °C for 5 min was performed, the samples were stored at 4 °C and triplicates were combined before gel electrophoresis on 1.5% agarose gels.

The PCR products were then normalized using the Invitrogen SequalPrep Plate Normalization Kit (Thermo Fisher Scientific), purified with AMPure beads (Agilent, Santa Clara, USA) and quality checked using High Sensitivity DNA Chips on a Bioanalyzer 2200 (Agilent). Before sequencing, the DNA pool was also quantified on a Qubit II Fluorometer using the dsDNA High-Sensitivity Assay Kit (Thermo Fisher Scientific). The final library pool was then loaded into a V2 2×250 cycle reagent Miseq cartridge according to the manufacturers protocol (Illumina, 2013, 2017) and sequenced on an Illumina Miseq device (Illumina Inc., San Diego, USA) at the Department of Human Genetics of the University of Würzburg, Germany. To account for low sequence diversity of the 16S rRNA library, an Illumina 5% PhiXv3 control library was added to the sequencing pool.

### Sequence Analysis

We used USEARCH v11.0.667 for the complete sequence analysis (Edgar, 2010). FASTQ sequences data were merged and length truncated with a minimum read length of 250 bp and maximum sequence differences of 10 base pairs of each sequence. After quality filtering, dereplication, singleton exclusion and chimera removal, sequences were denoised and dereplicated into amplicon sequence variants (ASVs) using the Unoise3 algorithm (Edgar, 2016a). The following taxonomy assignment was executed based on the RDP v16 reference database using bootstrap levels of 0.8 (Edgar, 2016b). A phylogenetic tree was constructed using FastTree 2.1.3 (Price et al., 2010). Further data analysis was conducted in R v3.5.2 (R Foundation, Vienna, Austria) using the package “phyloseq” (McMurdie & Holmes, 2013). ASVs that were assigned as chloroplasts or mitochondria and the 10 most abundant ASVs that were present in the negative controls were filtered from the dataset. All samples that had less than 1000 reads, were excluded from further analysis. The related raw data was deposited on the BExIS database of the Biodiversity Exploratories (www.bexis.uni-jena.de) with the dataset ID 26248.

### Analyzing Diversity of Bacterial Communities

To visualize and test for differences between the groups (species, tissue), a detrended correspondence analysis (DCA) and environmental fitting was performed using the R-package “vegan” (Oksanen et al., 2007). Functions of the “vegan” and “phyloseq” packages were further used to estimate alpha-diversity (Shannon, Richness) and beta-diversity indices (unweighted UniFrac, weighted UniFrac and Bray-Curtis dissimilarity). The “picante” package (Kembel et al., 2010) was used to calculate Faiths’ phylogenetic diversity (PD, Faith, 1992). Wilcoxon tests were performed on alpha- and beta-diversity values to test for statistically significant differences between plant species and plant organs. To determine ubiquitous microbiota (consistent ASVs among tissues), we used the R-package “microbiome”. As a detailed analysis for taxa of interest (Lactobacillales: association with pollinators; Rhizobiales: root bacteria), heatmaps were constructed and differences in relative abundance between groups were tested with t-tests. To identify chosen individual ASVs at the species level, 16S rDNA sequences were compared with available rDNA sequences in GenBank, using the NCBI BLASTN program (Zhang et al., 2000).

### Analyzing Biogeographic Differences

To test for differences in community composition between the three long distant bioregions, we applied environmental fitting models. The effect of distance on the shaping of the tissue specific microbial structures was further investigated through Mantel correlation between Bray-Curtis dissimilarities and geographical distances between each sample and sampling site. We applied ANOVA tests on bacterial classes for each tissue type to identify those representatives that were significantly different distributed between bioregions. Furthermore, we tested the potential turnover between tissue specific bacterial communities by comparing samples originated from same plots and those from different plots against each other, using pairwise Wilcoxon tests on Bray-Curtis dissimilarities.

### Analyzing Land use intensity effects

Measures of phylogenetic diversity were tested to identify differences between low and high LUI categories using Wilcoxon tests. To account for land use intensity within each of the groups, we computed permutational analysis of variance (ADONIS) tests for LUI categories on weighted and unweighted UniFrac distances and Bray-Curtis dissimilarities. The dispersion of samples within a LUI category (within group similarity) was estimated using the betadisper function. Differences between LUI categories were further tested on relative abundances of bacterial families using t-tests and Pearson correlations between bacterial taxa including environmental factors were analyzed using the R-package “psych” (Revelle, 2017).

All statistical tests above were tested for normal distribution and if they could not meet the requirements, their nonparametric test equivalents were analyzed. The confidence intervals were set at 95 % and p-values were adjusted for multiple testing with Benjamini-Hochberg correction (Benjamini & Hochberg, 1995). All visualizations were performed in R using the package “ggplot2” (Ginestet, 2011).

## Results

### Diversity of flower and leaf microbiomes

Illumina sequencing of the 16S rRNA gene region resulted in an average of 22,204 high-quality reads per sample of 406 samples in total after quality filtering. Accumulated over all samples of both plant species and plant organs, we found 5222 distinct ASVs.

Taxonomic classification of the microbial composition revealed 25 microbial phyla, including 4 phyla belonging to very low abundant archaeal taxa. All four tissues were dominated by Proteobacteria with numbers ranging from *R. acris* flowers (FRA 75 %), *T. pratense* flowers (FTP 90 %), *R. acris* leaves (LRA 77 %) to *T. pratense* leaves (LTP 63 %). Besides Proteobacteria, Bacteroidetes (FRA 9 %; FTP 6 %; LRA 29 %; LTP 11 %), Actinobacteria (FRA 1 %; FTP 1 %; LRA 6 %; LTP 6 %) and Firmicutes (FRA 8 %; FTP 3 %; LRA 1 %; LTP 3 %) were also abundant in all categories. However, each tissue type differed in their compositional structure, especially between plant organs (Supplementary Table S1). Interestingly, there were significantly different occurrences of Lactobacillales and Rhizobiales between flower and leaf samples (t-test, p < 0.001***). ASVs belonging to Lactobacillales were found in higher relative abundances on flowers than on leaves, while Rhizobiales were found more often on leaves (Supplementary Figure S1). In addition, all four tissue types shared ubiquitous bacteria (ASVs present across 95 % of all samples) all belonging to Proteobacteria with tissue specific differences in their relative abundances as shown in the Supplementary Table S2.

The number of ASVs and the diversity (Shannon’s index and Faith’s phylogenetic diversity) were generally significantly higher for leaves than flowers (Table 1). Evenness, an index for homogeneity in the distribution of ASV abundance, was also higher for leaves than flowers, indicating leaf communities were more homogenous with fewer highly abundant ASVs than those of flowers (p-value <0.001***, Table 1). Detrended correspondence analysis (DCA) showed separated clustering of bacterial communities by plant species and organs (Figure 1). This was confirmed by the environmental fitting model, with both plant species across organs (r2 = 0.29, p < 0.001***) and plant organs across species (r2 = 0.57, p < 0.001***) being statistically significant as explanatory factors and explaining large proportions of the overall community variance. Since the four different types of tissues showed strong differences, all following analyses were conducted separately within each sample type. Bray-Curtis dissimilarity analysis (multivariate homogeneity of group dispersions, F = 132.48, p < 0.001***) showed higher beta diversity in floral communities of *R. acris* (distance to centroid = 0.59±0.08) than in *T. pratense* (distance to centroid = 0.45±0.12). In leaf tissues, we observed a contrary pattern (multivariate homogeneity of group dispersions, F = 8.79, p < 0.005**). Communities of *R. acris* (distance to centroid = 0.45±0.12) were significantly less heterogeneous than leaf communities of *T. pratense* (distance to centroid = 0.50±0.11).

**Table 1:** Richness and diversity of the microbiome of different plant tissues. The given values represent averages and their respective standard deviations. Samples labelled with different letters were significantly different (analysis of variance difference of means with posthoc Tukey’s test, p < 0.05).

| Plant tissue | ASV Richness (S) | Shannon (H') | Evenness (H/logS) | Phylogenetic Diversity (PD) |
| --- | --- | --- | --- | --- |
| Flowers *R. acris* | 304.61±179.08 ab | 2.69±0.96 a | 0.48±0.16 a | 30.29±9.71 a |
| Flowers *T. pratense* | 271.42±83.82 a | 2.55±0.72 a | 0.46±0.12 a | 25.38±5.56 b |
| Leaves *R. acris* | 471.74±236.47 c | 3.82±0.58 b | 0.64±0.09 b | 34.25±10.54 c |
| Leaves *T. pratense* | 342.69±157.42 b | 3.75±0.81 b | 0.65±0.14 b | 29.98±8.66 a |

**Figure 1:**
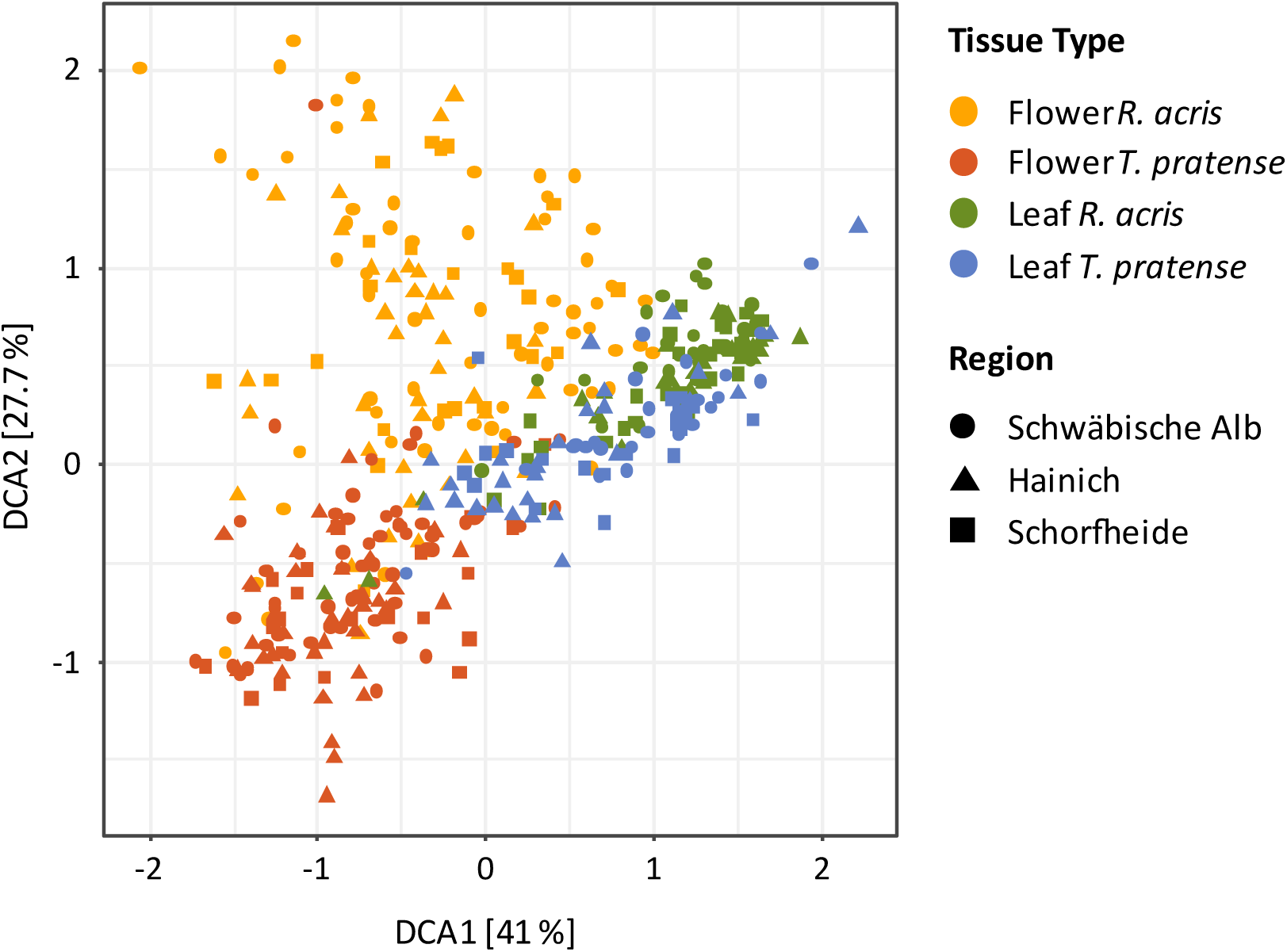
Ordination of flower and leaf microbiota samples of *R. acris* and *T. pratense* was conducted using detrended correspondence analysis (DCA) based on Bray-Curtis dissimilarity matrix. The calculation of beta diversity dissimilarities was performed on relative abundance of the full dataset that was not abundance filtered. Tissue sample types are colored differently and shaped according to their regional origin. Statistically significant differences were calculated by environmental fitting model between plant species (r2 = 0.29, p < 0.001***) and plant organs (r2 = 0.57, p < 0.001***).

### Biogeographic effects

To test for regional differences, we used environmental fitting models on DCA ordinations of each tissue specific community structure. With an exception for flower communities of *R. acris* (r^2^ = 0.09, p < 0.005**), we did not find significant differences between the three geographically distinct regions of the Biodiversity Exploratories (Figure 1).

In addition, we tested the correlation between Bray-Curtis dissimilarities and geographical distances for all samples and sampling sites of each tissue type. Here we confirmed our previous findings that bacterial communities of distinct bioregions with at least 300 km distance between each other had no separated structures. Neither *R. acris* flowers (Mantel test, r = −0.07678, p = 0.996) and *R. acris* leaves (Mantel test, r = −0.1043, p = 0.998) nor *T. pratense* flowers (Mantel test, r = −0.0896, p = 0.990) and *T. pratense* leaves (Mantel test, r = 0.006849, p = 0.634) were statistically significant correlated with distance. To identify differences in community compositions between distinct exploratory regions, we used ANOVA tests, which revealed that most abundant ASVs (filtered for min relative abundance 0.1 %) in the individual tissues were present in all three regions. Only ASVs linked to the taxonomic classes Cytophagia, Flavobacteriia, Bacilli and unclassified Proteobacteria showed significantly different occurrences within distinct bioregions (Supplementary Table S3). Taking these results together, bioregions can be considered to hold a very similar overall pool of flower and leaf bacteria even over long geographic distances.

Despite that, we still observed variability and ASV turnover between individual plants (Bray-Curtis dissimilarity ranges, Flowers *R. acris*, 0.74±0.2–0.83±0.04, Flowers *T. pratense*, 0.57±0.11–0.63±0.05, Leaves *R. acris*, 0.53±0.12–0.63±0.06, Leaves *T. pratense*, 0.55±0.13–0.70±0.06). Within bioregions, the turnover between two individuals was stronger if they originated from different plots than from the same plot (Wilcoxon signed rank test, Flowers *R. acris*, r = 0.69, p < 0.001***; Flowers *T. pratense*, r = 0.58, p < 0.001***; Leaves *R. acris*, r = 0.79, p < 0.001***; Leaves *T. pratense*, r = 0.84, p < 0.001***, Figure 2). This indicates that the local environment can affect microbiome composition already over short distances.

**Figure 2:**
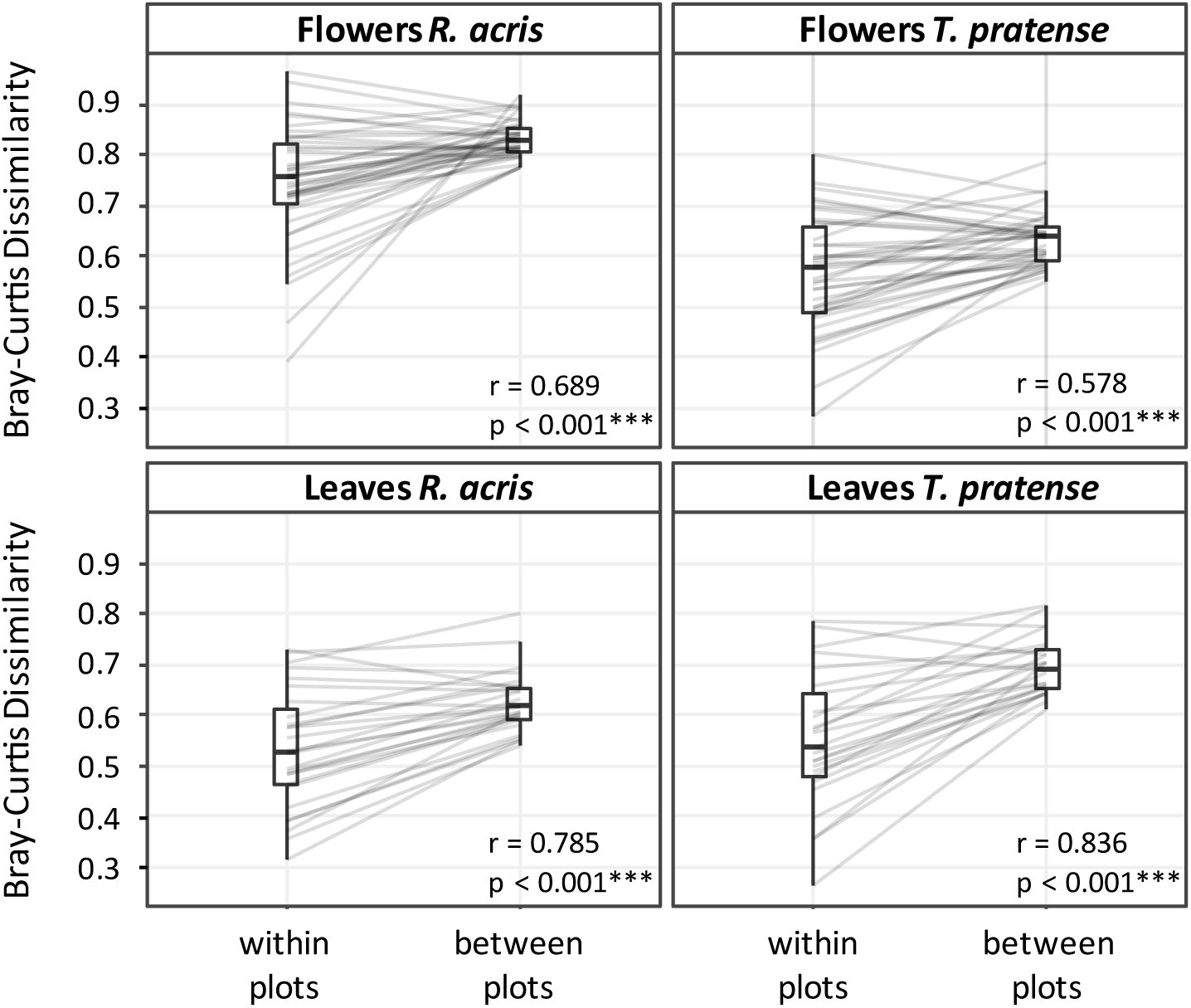
Tissue specific data showing mean beta-diversity between individuals within and between plots in the same bioregion. The boxplots indicate the first and third quartile and the median is displayed as horizontal line. Significant differences were tested using Wilcoxon signed rank tests.

### The impact of land use intensity

Land use intensity (LUI) affected the phylogenetic alpha-diversity (PD) significantly on flowers and leaves of *R. acris* and on leaves of *T. pratense* (Mann-Whitney-U test, p < 0.05*) with lower diversity, corresponding to less bacterial families, under high LUI, but not in flowers of *T. pratense* (Figure 3). However, the analysis of Shannon diversity revealed no significant changes caused by LUI in any of the investigated tissues (t-test, p > 0.05). Thus, while the species diversity remained mostly consistent, on higher phylogenetic levels less taxonomic groups were represented in microbiomes under high-LUI scenarios.

**Figure 3:**
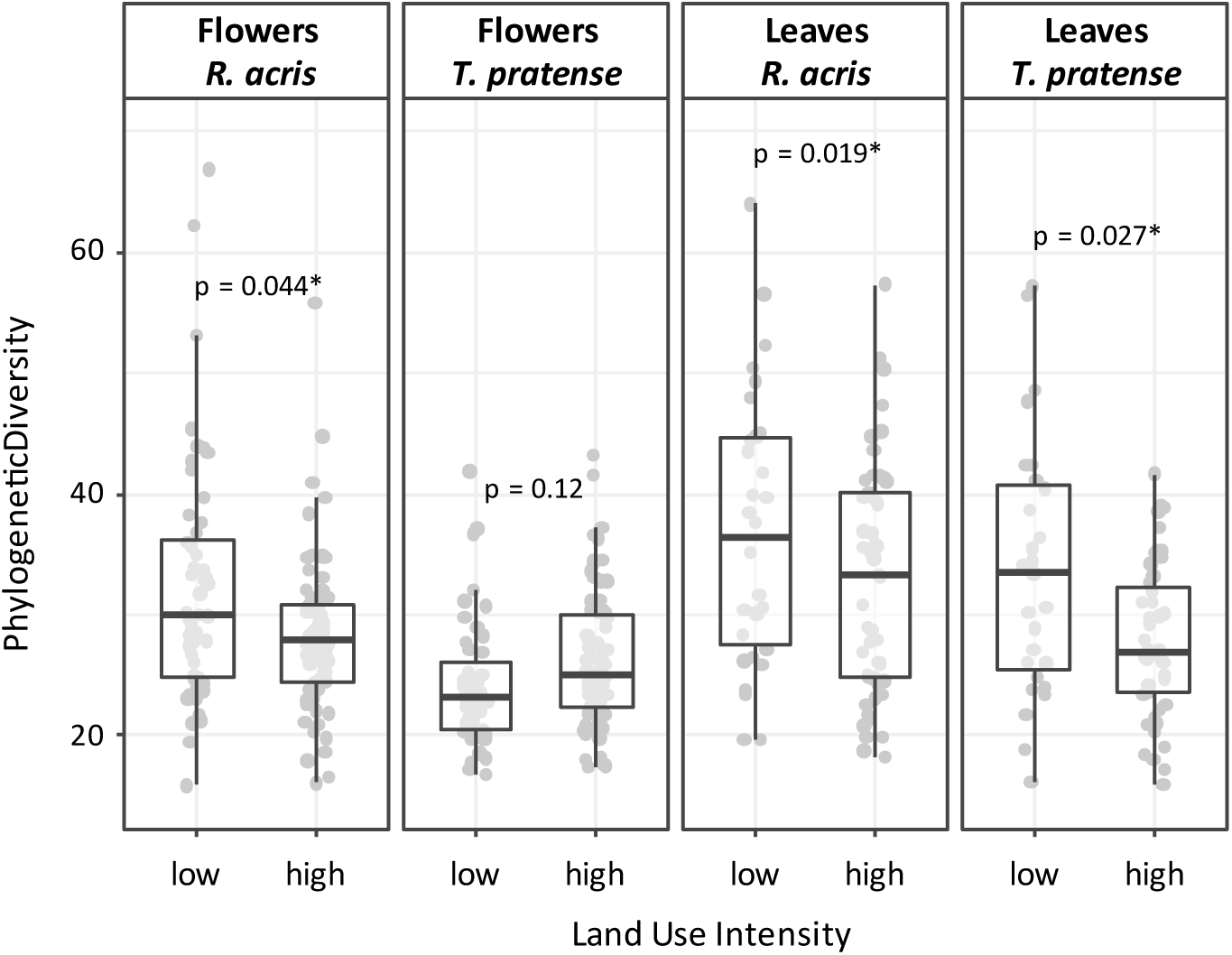
Faith’s phylogenetic diversity of all tissue types affected by low and high land use intensity managements regimes, respectively. The boxplots indicate the first and third quartile and the median is displayed as horizontal line. Significant differences between low and high LUI are indicated with an asterisk (Mann-Whitney-U test).

To test for changes in community structures caused by LUI, we analysed weighted (absence/presence, abundance, phylogeny) and unweighted UniFrac distances (absence/presence, phylogeny) as well as Bray-Curtis dissimilarities (absence/presence, abundance). Significant differences in beta-diversity between low and high LUI were detected on unweighted UniFrac distances in all tissues using permutational multivariate analysis of variance (ADONIS) (Supplementary Table S4). Analysis of Bray-Curtis dissimilarity and weighted UniFrac distance revealed no significant changes in response to LUI. Therefore, significant changes of bacterial communities were driven rather by qualitative patterns (presence or absence of certain ASVs) than by abundance-based dissimilarities.

To identify separately the effects of each land use management, we tested changes in unweighted-UniFrac distances of each tissue type against grazing, mowing and fertilization intensity categories (Figure 5). Flower and leaf microbiomes of all tissues (multivariate analysis of group dispersions, Flowers *R. acris*, F = 3.8, p = 0.012*; Flowers *T. pratense*, F = 3.6, p = 0.015*, Leaves *R. acris*, F = 4.8, p = 0.007**, Leaves *T. pratense*, F = 13.4, p < 0.001***) were more homogenous in very intensively mown landscapes than in less intensively managed fields, as revealed by tukey’s post-hoc test of betadisper results. High fertilization intensity significantly led to lower beta diversity of community structures on *R. acris* flowers (F = 2.9, p = 0.041*) and *R. acris* leaves (F = 3.7, p = 0.015*) as well as on *T. pratense* leaves (F = 6.8, p = 0.002**), but not on flowers of *T. pratense* (F = 1.5, p = 0.206). Furthermore, very intensive grazing management had only minor effects on floral communities of *T. pratense* (F = 3.7, p = 0.015*), but did not change the heterogeneity of bacterial community structures of other tissue types (p > 0.05). This suggests that the different types of land use manipulate the microbial community structures in different ways.

**Figure 4:**
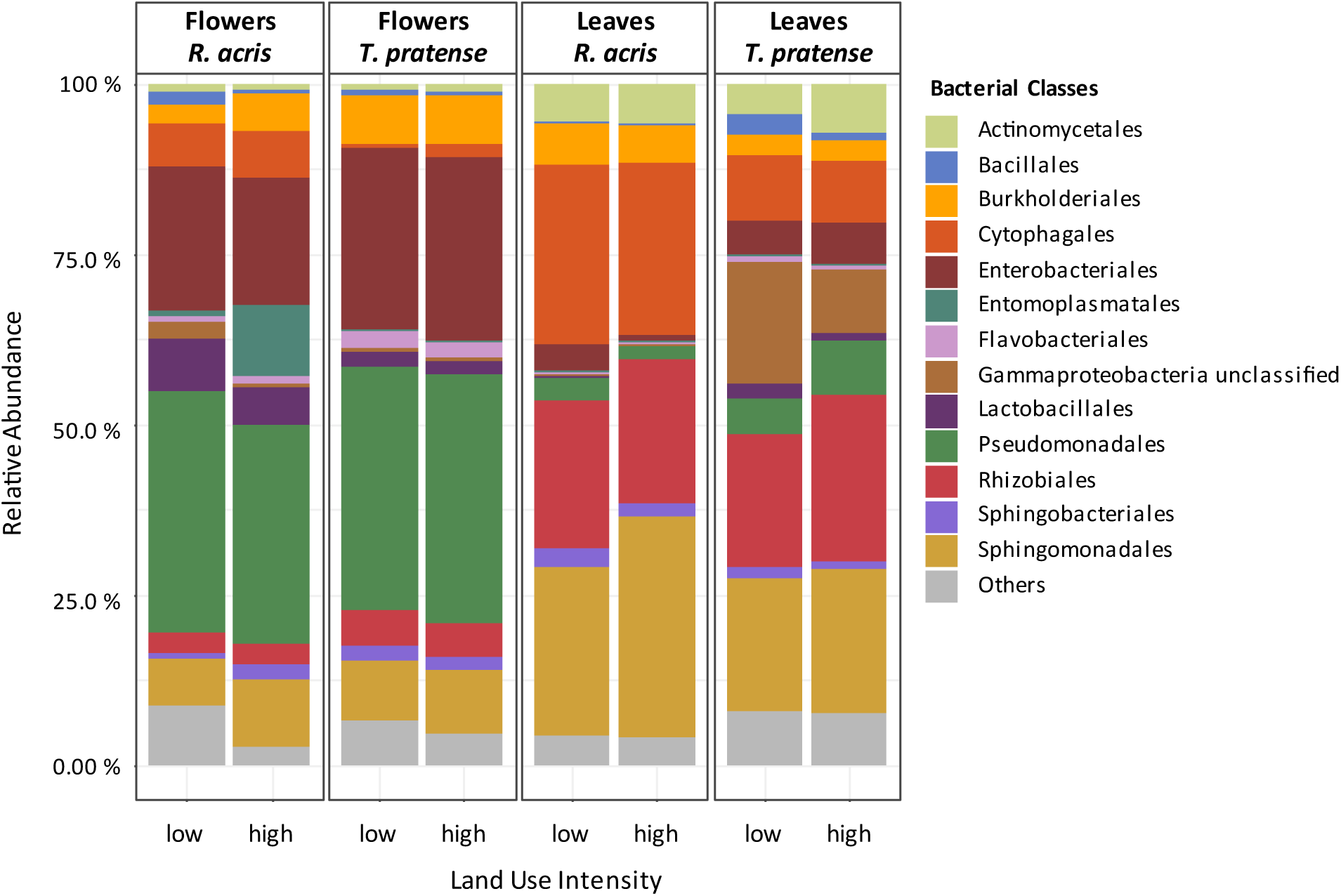
Bacterial composition of all four tissue types in low and high land use intensity environments. Relative abundance represents the mean distribution in 16S rDNA sequence reads over samples and tissue types. Microbial orders 1 relative abundance were grouped as “Others”.

**Figure 5:**
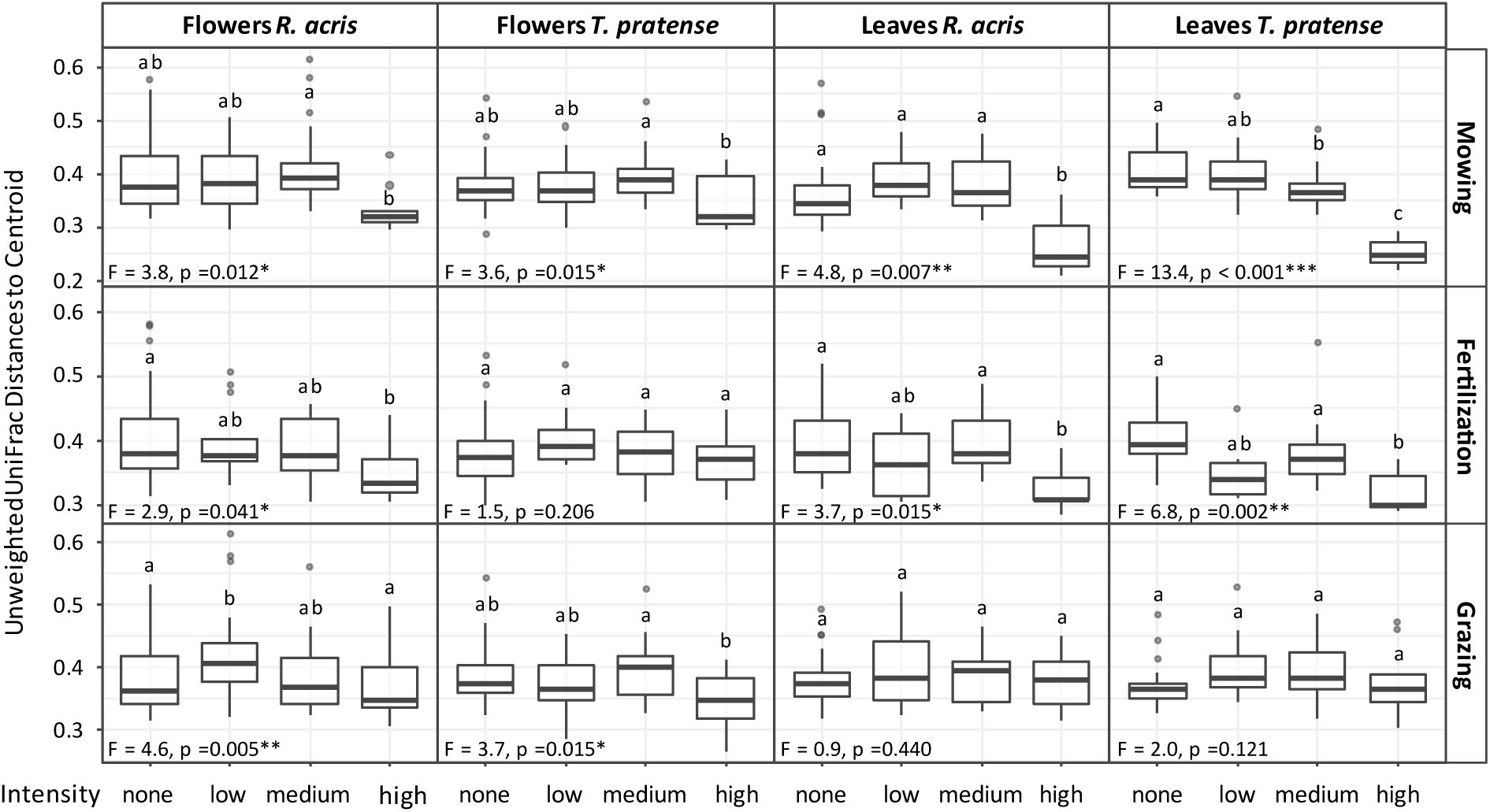
The impact of mowing, fertilization and grazing intensities on variabilities in community structures of different plant species and tissue types. The analysis of unweighted UniFrac represent the beta-diversity as distances to group centroids of each community. Differences between the intensity categories of each land use management type were assessed by multivariate analysis of group dispersions (betadisper). Different letters indicate statistical differences between these categories as revealed by Tukey’s post-hoc test (p < 0.05), respectively for each land use management and each plant species and organ. Effects of high land use managements were revealed for mowing and fertilization for almost all tissue types. Grazing had only minor consequences on the bacterial community structures.

The bacterial compositions of each tissue in low and high LUIs did not differ much from each other (Figure 4). Moreover, it seems like all flower and leaf organs have been constant in variable land use regimes. However, there were changes in relative abundance for some bacterial taxa, which were specific for plant tissues (Supplementary Tables S5A-D). For example, on flowers of *R. acris* there was an increase of Entomoplasmatacae (Entomoplasmatales) under high LUI (p = 0.047*, mean relative abundance, low LUI = 0.79 %, high LUI = 6.16 %). The family of Entomoplasmataceae was represented by 5 ASVs including the most abundant species *Mesoplasma florum* as NCBI BLASTN best scores revealed. The genus *Mesoplasm*a showed increased relative abundance with higher mowing and fertilization intensities (Supplementary Figure S2A). Overall, a striking pattern was that in all four tissues always one or two abundant bacterial families were either negatively or positively affected by high LUIs, while a variety of other bacterial families were always less abundant in the opposing LUI environment (Supplementary Tables S5A-D). Therefore, it is likely that especially the former benefit from the given environmental conditions.

Because our results revealed that some bacterial families were affected by LUI, we w.anted to have deeper insights whether bacterial genera were correlated with land use types. Investigating the 25 most abundant genera, we found that not many taxa were correlated with LUI (Supplementary Figures S2A-D). However, we identified bacteria of the genus *Spirosoma* on *R. acris* flowers and on *T. pratense* leaves occurring more abundant under higher grazing. Furthermore, we found on leaves of *T. pratense* and *R. acris* positive correlations between grazing and some Actinobacteria as well as *Chryseobacterium*, suggesting that there might be mechanisms of this land use type that support the establishment of these taxa.

## Discussion

Bacterial colonizers of above ground-plant parts are thought to be affected by a variety of plant-based and environmental characteristics, which are so far not fully understood. Especially the diversity and community composition of floral bacteria were poorly considered with respective to their environmental determinants. In our study, we therefore investigated the factors plant species and plant tissue identity as well as biogeography and land use management that revealed different effects on the structural organization of bacterial communities.

### Diversity of flower and leaf microbiomes

The analysis of alpha-diversity indices (Shannon and PD) showed that leaves harbour more diverse microbiomes than flowers, which has also been reported in other studies (Junker & Keller, 2015; Junker et al., 2011; Krimm et al., 2005; Leff et al., 2015). A possible explanation for the higher diversity on leaf surfaces could be due to unspecific availability of a variety of resources including carbohydrates, amino acids, fatty acids, sugars and alcohols that can be consumed by many bacterial taxa, while flowers potentially prohibit the establishment of various bacteria due to their specific biochemical profile (Vorholt, 2012). This exclusion of several bacteria is likely due to the sugar-rich secretions of floral structures like nectaries. Thus, only bacteria that are able to adapt to this osmotic stress are able to maintain within the given environment (Aleklett et al., 2014). In addition, flowers emit specific volatile compounds that could inhibit the growth of bacteria, that leaves do not produce, as suggested by Junker and Tholl (2013). Consequently, bacteria that are also able to resist against plant defense mechanisms as antimicrobial compounds of nectar would be favoured and therefore potentially found in higher proportions. Altogether, these assumptions might explain why we also found more evenly distributed communities on leaves than on flowers.

In our study, the overall microbial structures were found to be significantly distinct between plant organs and plant species, with plant organs accounting for greater variance. Other studies on different organs of specific plant species (Junker & Keller, 2015; Wei & Ashman, 2018; Zarraonaindia et al., 2015) and on bacterial leaf communities between different plant species (Kembel et al., 2014; M. Kim et al., 2012; Laforest-Lapointe et al., 2016; Lambais et al., 2006) confirmed these findings, suggesting organ specificity as the more important driver of overall bacterial community structures. This is also in line with a recently published study on *Ranunculus acris, Trifolium pratense* and *Holcus lanatus* of Massoni et al. (2019). Nevertheless, they also reported that both plant species and plant organs share almost all detected taxa when taking only absence and presence into account, which suggests exchange between tissues or environmental spill-over. Accordingly, we also found higher abundant taxa shared between both plant species and tissue types. Therefore, it is assumed that general taxa of phyllosphere bacteria can be ubiquitous among different plant species and organs, even though their proportional occurrence may vary strongly as follows.

While the tissues of both plant species were dominated by Proteobacteria, their relative composition of lower taxa levels differed fundamentally, especially between plant organs. On both plant species we found typical bacterial colonizers that have been reported in previous studies about flower and leaf microbiota (Bulgarelli et al., 2013; Junker & Keller, 2015; Junker et al., 2011; Krimm et al., 2005; Ottesen et al., 2013; Steven et al., 2018; Vorholt, 2012). Flowers were dominated by Pseudomonas, *Enterobacteriaceae* and *Sphingomonas*, varying in their relative abundances on each plant species. The most abundant bacterial taxa on leaves were represented by *Sphingomonas, Methylobacterium* and *Hymenobacter* with different frequencies per plant species. Members of these taxa might be of great importance for defending host plants against pathogens, passively by building a biological barrier on the surface of flowers and leaves (biological control agents) or actively by producing antimicrobial compounds (Innerebner et al., 2011; Volksch & May, 2001; Wilson & Lindow, 1993). The most abundant taxa found on flowers *Pseudomonas sp*., *Pantoea agglomerans* (Enterobacteriaceae) and *Pseudomonas syringae* are known for their plant growth promoting potential (Preston, 2004; Shariati et al., 2017). Interestingly, it is assumed that *P. syringae* and *P. agglomerans* are also involved in modifying floral chemistry and pollinator behaviour, thus affecting the success of host reproduction (Farré-Armengol & Junker, 2019).

We further found that members of Lactobacillales were solely and consistently present in higher proportions on flowers. Both plant species are visited by different insects, which supports the hypotheses, that these bacteria are likely introduced by pollinators (McFrederick et al., 2013; McFrederick & Rehan, 2019; McFrederick et al., 2017; Voulgari-Kokota et al., 2019c; Voulgari-Kokota et al., 2019b). On the other hand, Rhizobiales, especially representatives of the genus Methylobacterium, were found in higher relative abundances in leaf samples. Methylobacteria are able to utilize plant-derived compounds like methanol and methylamine as well as C2, C3 and C4 compounds as carbon sources and secreting plant growth promoting cytokinins, potentially favouring these bacteria to limited resources on the surface of leaves (Delmotte et al., 2009).

Bacteria that are associated with the upper surface of flowers and leaves need to be well adapted to these microhabitats. For example, they have to be protected against high ultraviolet radiation and must be able to endure reactive oxygen species or persist drought, temperature changes, water and osmotic stress. Especially extracellular polysaccharides (EPS), but also lipopolysaccharides (LPS), lipids and carotenoids are thought to be the major protective factors that bacteria produce to form aggregates and to overcome these stresses (Aragon et al., 2017; Lindow & Brandl, 2003; Schlechter et al., 2019; Stone et al., 2018; Vorholt, 2012). Especially taxa such as *Pseudomonas*, Enterobacteriaceae, *Sphingomonas, Methylobacterium* and *Hymenobacter* are able to produce a variety of these compounds (Danhorn & Fuqua, 2007; C. E. Morris & Monier, 2003) and consequently, these taxa could be found more likely and in higher abundances on these surfaces, even though this linkage has not been investigated experimentally so far. Moreover, it is supposed that the acquisition of bacteria from different environmental sources and their subsequent maintenance potential is limited, mainly by plant based factors (Rebolleda-Gómez et al., 2019; Vorholt, 2012) or bacterial competition (Hibbing et al., 2010). In addition, it is also suggested that the heterogeneity of pollinators that potentially transmit bacteria to flowers greatly contribute to microbial structures. This is especially supported by our findings of higher beta diversity on flowers of *R. acris* that are visited by 54 different insect species, especially syrphids and bees (Steinbach & Gottsberger, 1995), while flowers of *T. pratense* that are mainly visited by only bumblebees and honey bees (Free, 1993) showed less variability. It was reported that bacterial beta diversity was higher in nectar with increased heterogeneity of pollinator visitation (Vannette & Fukami, 2017) or in seeds that were not excluded to bee visitation (Prado et al., 2019).

Altogether, the differences in diversity, structure and composition between plant species and tissues could be explained by different surface structures with specific chemical compounds or by different ecological characteristics like exposure time within environments or pollinator variations. We thus investigated for further analysis each tissue type separately.

### Biogeography

The overall microbial structures did not differ between exploratory regions. However, we found that bacterial turnover was higher between individuals from different sites of the same region than individuals collected at the same site. These results may suggest that in addition to pheno- and chemotype, determinative characteristics of the very local environment could be of more importance than regional differences. This supports findings of previous studies with tissue specificity on *Metrosideros polymorpha* (Junker & Keller, 2015) and on *Cycas panzhihuaensis* (Zheng & Gong, 2019), or in another study on different Agave plants (Coleman-Derr et al., 2016), in which bacterial communities differed significantly between plant compartments or species, but not much between distinct bioregions. Furthermore, it was reported that geographic distance of bacterial communities on leaves of *Pinus ponderosa* trees had little influence with more variation at individual sites than between trees located on different continents (Redford et al., 2010). This contributes to the assumption that the previously described plant-based characteristics together with local environmental conditions are of major importance driving the assembly of bacterial communities.

### Land use intensity

We observed a decrease in phylogenetic diversity and number of bacterial families under intensive land use in both species and organs, with exception of the *T. pratense* flower microbiome. These findings correspond to another study on *Dactylis glomerata* root-associated microbes (Estendorfer et al., 2017) and might be due to selective processes that affect bacteria as a consequence of environmental disturbance. The reduced phylogenetic variance could indicate that its functional diversity is probably also limited because bacterial species traits often reflect shared evolutionary history (Mazel et al., 2018; but see Aminov, 2011; Burke et al., 2011 or Milner et al., 2019). It is supposed that only bacteria with adaptation or recolonization strategies to frequently altered environments could be found within these plant microhabitats (Shade et al., 2012). Contributing to this, those taxa that were also dominant in low LUI environments, were not negatively affected by land use intensification. This is likely attributed to the fact that many phyllosphere bacteria recover or recolonize relatively quickly after disturbance events, which might be promoted by their ubiquitous lifestyle and by their fast reproduction rates. The fact that the highly abundant genera can be found in several habitats could be of great importance for their reestablishment. The genera *Pseudomonas, Pantoea, Sphingomonas, Methylobacterium and Hymenobacter* constitute a considerable and generally stable fraction of phyllosphere bacterial communities of terrestrial plants under varying environmental conditions (Aleklett et al., 2014; Knief et al., 2010; Lindow & Brandl, 2003). The adaptation to these microhabitats and resistance or resilience against environmental changes might be promoted by these dominant genera modifying their immediate environment (Schlechter et al., 2019).

Furthermore, we found that especially very intensive mowing and fertilization led to more homogeneous communities. The differences were rather based on qualitative patterns than abundances. Thus, it is supposed that these changes on bacterial community structures are mainly attributed to low-abundant taxa. Remarkably, Pearson correlations revealed autocorrelations between the three LUI types. Both mowing and fertilization were positively co-correlated, also with overall LUI, while grazing was negatively correlated with mowing and fertilization. Therefore, we assume that high LUI was mainly due to high mowing and fertilization rates, and both had consequences on reduced turnover of the flower and leaf community structures, while grazing had only minor effects. These synergistic effects of high mowing and grazing intensities were also reported in studies on bacterial endophytes of different grass species (Wemheuer et al., 2017; Wemheuer et al., 2016). They further showed that plant species identity was the main driver of bacterial assembly, but agronomic management practice had also an impact on their composition. Moreover, one of these studies revealed that land use intensification affected both the functionality and the taxonomy of bacterial communities, which, however, were not correlated with each other (Wemheuer et al., 2017). This indicates that a reduction in phylogenetic diversity is not necessarily related to a loss in functionality. Considering that the core microbiota is of functional great importance for the plant holobiont (Vandenkoornhuyse et al., 2015), which would support our findings of ubiquitous and highly abundant genera that were not affected by land use intensification.

In bacterial flower communities of *R. acris*, Entomoplasmatales were predominantly found in regularly disturbed environments, particularly with frequent mowing and high fertilization rates. This taxon is known to have very short replication times and would favour sugar rich environments. The most abundant member was referred to *Mesoplasma florum*, which has a very small genome size (∼ 800 KB), is assumed to be non-plant-pathogenic and known to be transmitted between flowers by insects (Baby et al., 2018). Due to their very high mutation rates, this species can rapidly adapt to their surroundings and to environmental changes, which could favour them against other bacteria (Denamur & Matic, 2006; Sung et al., 2012). We further found no significant correlation effect for grazing, which was likely since ungrazed plots had high mowing and fertilization rates. Keeping in mind that *R. acris* produces a toxic compound named protoanemonin, that can be recognizable by livestock, these plants were likely not grazed at all (Lamoureaux & Bourdot, 2007). It has its highest concentration at the flowering stage, especially when the plant is crushed, and shows antibacterial activity against a broad spectrum of bacteria by inhibiting quorum sensing (Bobadilla Fazzini et al., 2013; Didry et al., 1993). Thus, together with other antimicrobial compounds it has the potential to determine bacterial community compositions. This could contribute to the higher abundance of *Mesoplasma* and further, to our findings that bacterial communities of *R. acris* flowers and leaves were more homogenous in high LUI environments, when plants were exposed to increased environmental disturbance. Given these results, floral *Mesoplasma* was a strong disturbance indicator taxon for *Ranunculus* flowers.

Neither mowing, grazing nor fertilization affected the bacterial community on floral tissues of *T. pratense*, which could be due to the growth height of this species, which is relatively low in comparison to that of *R. acris*. Thus, flowers of that plant species suffered less from mowing (here at heights of mostly 7-8 cm). However, higher mowing intensity led to a reduction of beta-diversity on leaves of *T. pratense*, even though no bacterial genera were affected by this land use type. This supports our assumptions that only the minor abundant taxa were affected by land use and that the dominating bacteria are resilient and able to re-establish on the re-growing tissues. We further found that grazing did not affect the heterogeneity of bacterial communities. However, for leaves of both plant species on the other hand, our results revealed a positive correlation between the relative abundance of two genera related to Actinobacteria and grazing. A study on soil microbiota of Shange et al. (2012) found that Actinobacteria were more abundant in intensively grazed landscapes. One explanation for this could be, that Actinobacteria are generally more abundantly present in those environments or leaves particularly often exposed to soil bacteria when stamped down by cattle. Wagner et al. (2016) reported that roots and leaves of a wild mustard plant (*Boechera stricta*) shared highly similar bacterial communities supporting the assumption that bacterial community members could be recruited from the soil. In other studies on the rhizosphere or roots, Actinobacteria profited from abiotic stresses of host plants, especially from drought (Naylor & Coleman-Derr, 2018; Sathya et al., 2017). Two further bacterial genera were positively affected by grazing, *Spirosoma* found on leaves of *T. pratense*, but also on flowers of *R. acris* and *Chryseobacterium* on leaves of *R. acris*. These two genera were reported from the rumen of cows (Huws et al., 2015; Thomas et al., 2017). Thus, it is likely that they were introduced by grazing cattle.

Altogether, our results suggest that more frequently disturbed environments might favour highly competitive and fast-growing microbes that are resilient to land use intensification, yet this remains to be assessed in future studies. In addition, when we consider the bacterial structures of each individual tissue type, it seems they are rather determined by individual plant-based factors, most likely by surface morphology or biochemistry, than by anthropogenic changes of the environment.

## Conclusion

To our knowledge, here we provide a first study that examined bacterial community compositions of flowers and leaves with respect to biogeography and land use intensity. Our key findings included that bacterial communities: (1) were different between flowers and leaves and varied also between the two investigated plant species; (2) showed turnover between short-distance locations, which was not the case for larger distances between distinct bioregions (3) were less diverse and more homogenous in environments with high land use intensity, with no negative effects on those taxa that were even dominant at low intensities. Future studies and experiments on plant functional traits, especially by phytochemical properties, could prove highly interesting to understand the mechanisms behind this resilience. Additionally, the metabolic potential of the most dominant and ubiquitous phyllosphere bacterial members and their functional contribution to plant health and fitness can provide valuable insights in following studies.

## Acknowledgements

We thank the managers of the three Exploratories, Kirsten Reichel-Jung, Iris Steitz and Sandra Weithmann (Schwäbische Alb), Juliane Vogt (Hainich), Miriam Teuscher (Schorfheide), and all former managers for their work in maintaining the plot and project infrastructure; Christiane Fischer for giving support through the central office; Andreas Ostrowski for managing the central database; and Markus Fischer, Eduard Linsenmair, Dominik Hessenmöller, Daniel Prati, Ingo Schöning, François Buscot, Ernst-Detlef Schulze, Wolfgang W. Weisser, and the late Elisabeth Kalko for their role in setting up the Biodiversity Exploratories project. Fieldwork permits were issued by the responsible state environmental offices of Baden-Württemberg, Thüringen, and Brandenburg.

