## Supplemental figures and tables for "Changes amid constancy: flower and leaf microbiomes along land use gradients and between bioregions"

### Supplementary Figures and Tables

#### Figures

**Figure S1: Heatmap with relative abundance of Lactobacillales and Rhizobiales ASVs of each sample related to tissue type.** Differences in their occurrence on flowers and leaves (plant organs) were statistically tested using t-test ($p < 0.001$***).

**Figure S2A-D: Correlations between relative abundances of 25 most abundant bacterial genera and LUI parameters**. Correlations are based on linear Pearson correlation coefficients against each other and LUI indices. Correlation coefficients are displayed by the scale color in the filled squares and indicate the strength of the correlation (r) and whether it is positive (blue) or negative (red). P-values were adjusted for multiple testing with Benjamini-Hochberg correction and only significant correlations are shown ($p < 0.05$). White boxes indicate non-significant correlations. ***A)*** *Ranunculus acris* flowers, **B)** *Trifolium pratense* flowers, **C)** *Ranunculus acris* leaves (LRA), **D)** *Trifolium pratense* leaves (LTP).

#### Tables

**Table S1: Taxonomic identification of the most abundant bacterial genera and their presence** (average in percent) on each tissue type.

**Table S2: Taxonomic identification of ubiquitous bacteria found in 95 % of all samples**, including their average relative abundance on each tissue type.

**Table S3: Bacterial Classes that differed significantly in relative abundance between bioregions for each tissue type.** Differences between bioregions were tested using ANOVA. Relative abundances are shown for each exploratory region in percentage).

**Table S4: The impact of LUI on beta-diversity of bacterial community compositions of different plant tissues.** Statistical tests were performed using permutational multivariate analysis of variance (ADONIS) showing significance symbols ($p < 0.05$* and $p < 0.01$**).

**Table S5A-D: Bacterial families that were significantly affected by land use intensity.** Significant differences between high and low LUI were tested using t-test (confidence interval 95 %). P-values were adjusted for multiple testing with Benjamini-Hochberg correction. Bold numbers indicate higher values of mean relative abundances. **A)** *Ranunculus acris* flowers, **B)** *Trifolium pratense* flowers, **C)** *Ranunculus acris* leaves (LRA), **D)** *Trifolium pratense* leaves (LTP).

**Figure S1**

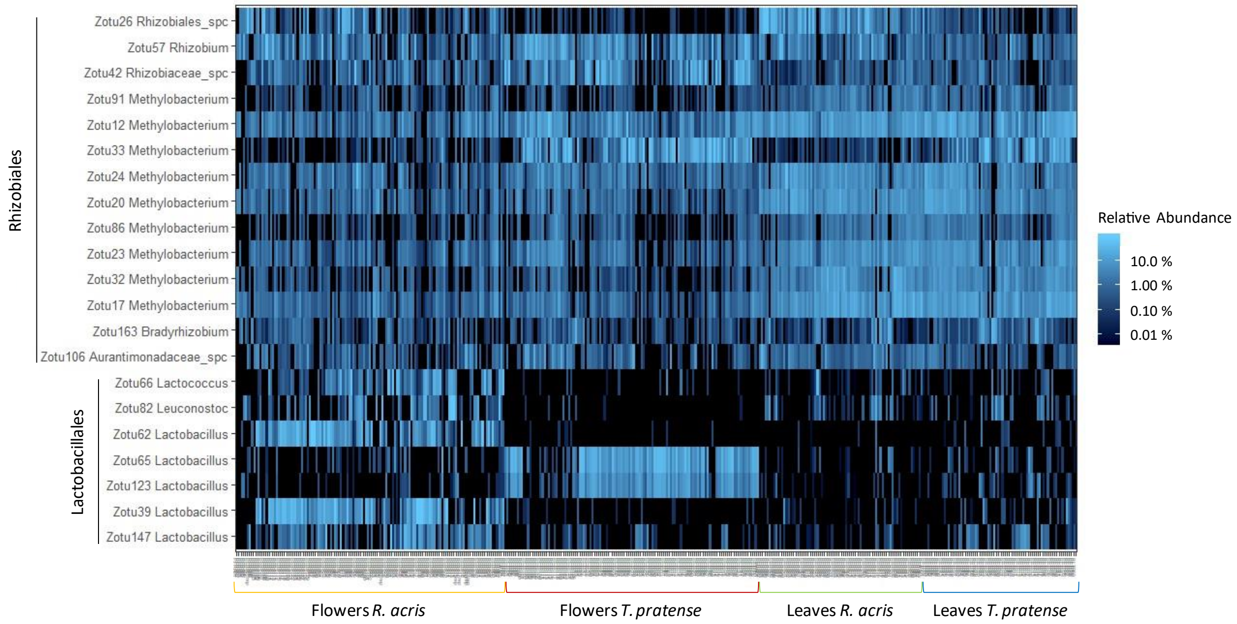

Rhizobiales
Lactobacillales
Zotu26 Rhizobiales_spc
Zotu57 Rhizobium
Zotu42 Rhizobiaceae_spc
Zotu91 Methylobacterium
Zotu12 Methylobacterium
Zotu33 Methylobacterium
Zotu24 Methylobacterium
Zotu20 Methylobacterium
Zotu86 Methylobacterium
Zotu23 Methylobacterium
Zotu32 Methylobacterium
Zotu17 Methylobacterium
Zotu163 Bradyrhizobium
Zotu106 Aurantimonadaceae_spc
Zotu66 Lactococcus
Zotu82 Leuconostoc
Zotu62 Lactobacillus
Zotu65 Lactobacillus
Zotu123 Lactobacillus
Zotu39 Lactobacillus
Zotu147 Lactobacillus
Relative Abundance
10.0 %
1.00 %
0.10 %
0.01 %
Flowers *R. acris*
Flowers *T. pratense*
Leaves *R. acris*
Leaves *T. pratense*

**Figure S2A**

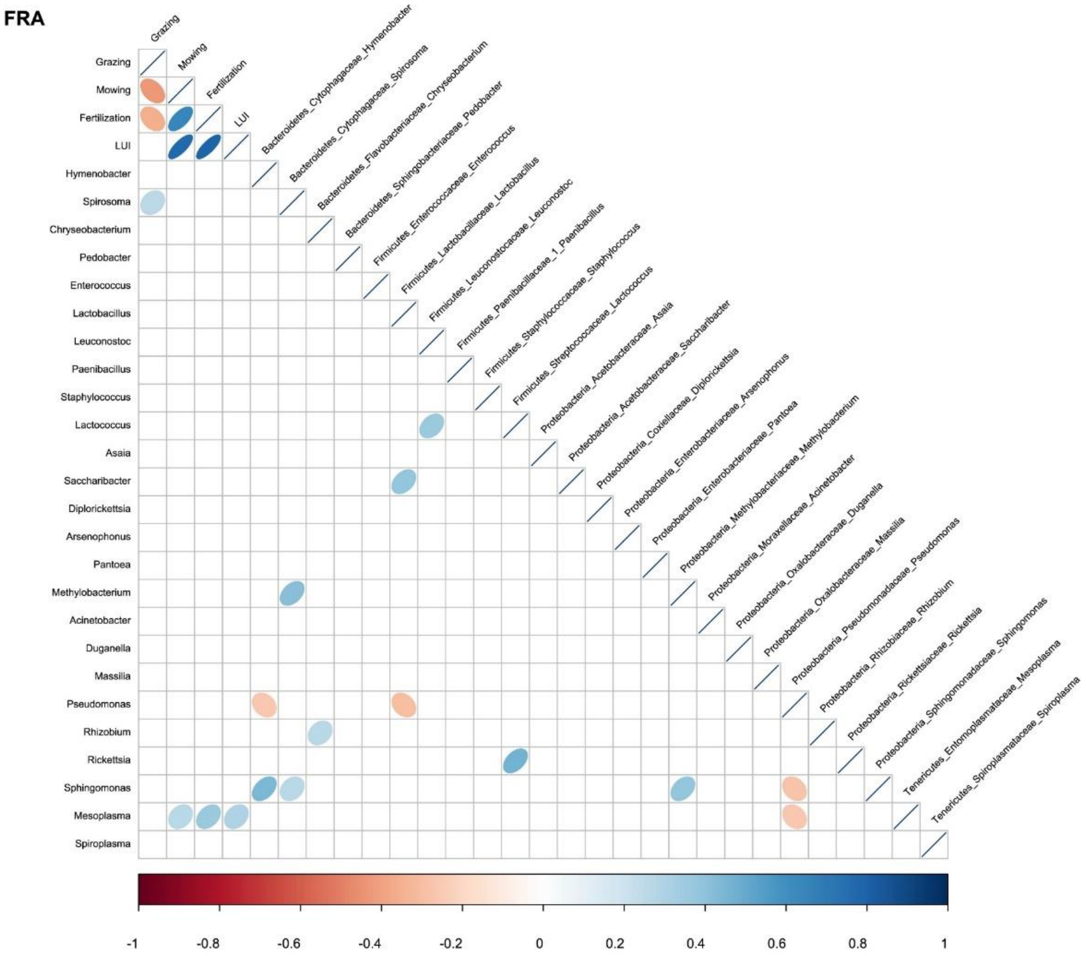
FRA
Grazing
Mowing
Fertilization
LUI
Bacteroidetes_Cytophagaceae_Hymenobacter
Bacteroidetes_Cytophagaceae_Spirosoma
Bacteroidetes_Flavobacteriaceae_Chryseobacterium
Bacteroidetes_Sphingobacteriaceae_Pedobacter
Firmicutes_Enterococcaceae_Enterococcus
Firmicutes_Lactobacillaceae_Lactobacillus
Firmicutes_Leuconostocaceae_Leuconostoc
Firmicutes_Paenibacillaceae_1_Paenibacillus
Firmicutes_Staphylococcaceae_Staphylococcus
Firmicutes_Streptococcaceae_Lactococcus
Proteobacteria_Acetobacteraceae_Asaia
Proteobacteria_Acetobacteraceae_Saccharibacter
Proteobacteria_Coxiellaceae_Diplorickettsia
Proteobacteria_Enterobacteriaceae_Arsenophonus
Proteobacteria_Enterobacteriaceae_Pantoea
Proteobacteria_Methylobacteriaceae_Methylobacterium
Proteobacteria_Moraxellaceae_Acinetobacter
Proteobacteria_Oxalobacteraceae_Duganella
Proteobacteria_Oxalobacteraceae_Massilia
Proteobacteria_Pseudomonadaceae_Pseudomonas
Proteobacteria_Rhizobiaceae_Rhizobium
Proteobacteria_Rickettsiaceae_Rickettsia
Proteobacteria_Sphingomonadaceae_Sphingomonas
Tenericutes_Entomoplasmataceae_Mesoplasma
Tenericutes_Spiroplasmataceae_Spiroplasma
Grazing
Mowing
Fertilization
LUI
Hymenobacter
Spirosoma
Chryseobacterium
Pedobacter
Enterococcus
Lactobacillus
Leuconostoc
Paenibacillus
Staphylococcus
Lactococcus
Asaia
Saccharibacter
Diplorickettsia
Arsenophonus
Pantoea
Methylobacterium
Acinetobacter
Duganella
Massilia
Pseudomonas
Rhizobium
Rickettsia
Sphingomonas
Mesoplasma
Spiroplasma
-1
-0.8
-0.6
-0.4
-0.2
0
0.2
0.4
0.6
0.8
1

**Figure S2B**

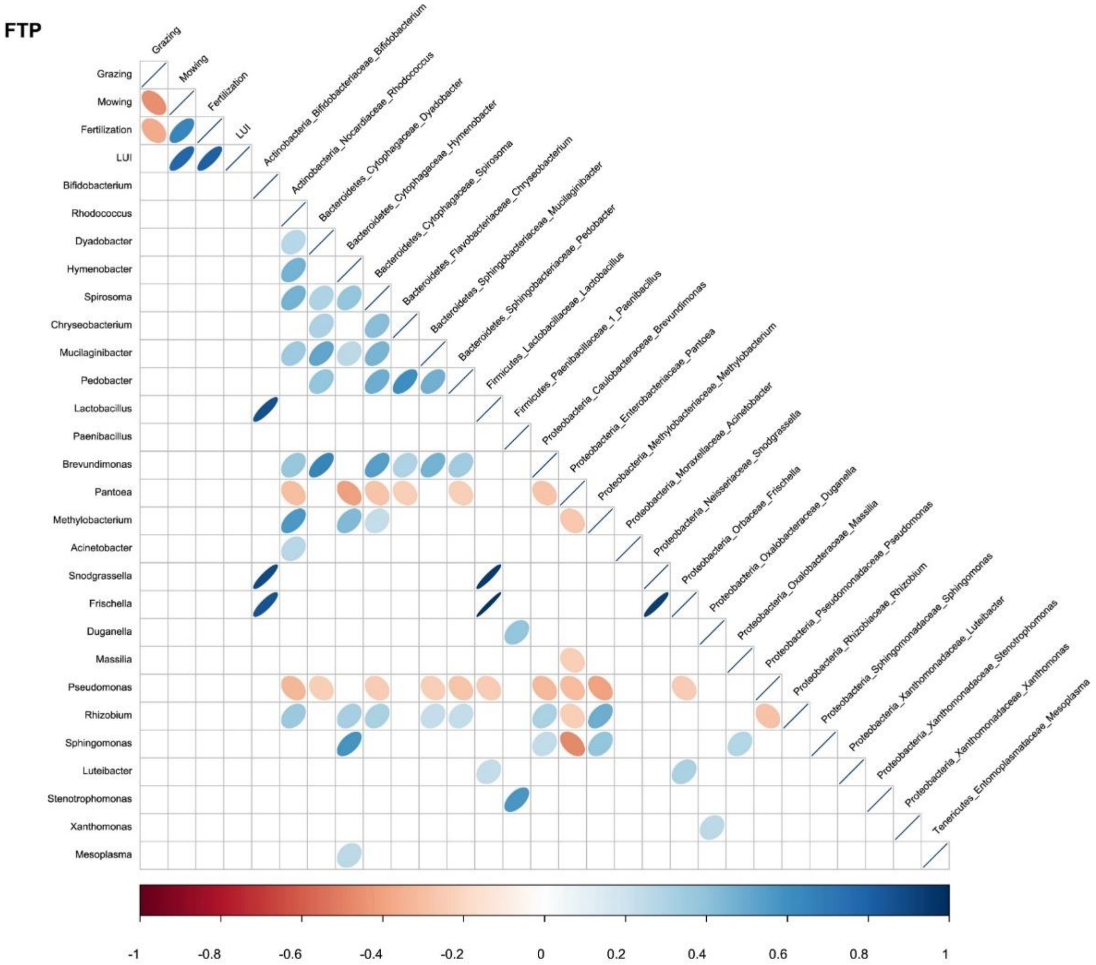
FTP
Grazing
Mowing
Fertilization
LUI
Bifidobacterium
Rhodococcus
Dyadobacter
Hymenobacter
Spirosoma
Chryseobacterium
Mucilaginibacter
Pedobacter
Lactobacillus
Paenibacillus
Brevundimonas
Pantoea
Methylobacterium
Acinetobacter
Snodgrassella
Frischella
Duganella
Massilia
Pseudomonas
Rhizobium
Sphingomonas
Luteibacter
Stenotrophomonas
Xanthomonas
Mesoplasma
Actinobacteria_Bifidobacteriaceae_Bifidobacterium
Actinobacteria_Nocardiaceae_Rhodococcus
Bacteroidetes_Cytophagaceae_Dyadobacter
Bacteroidetes_Cytophagaceae_Hymenobacter
Bacteroidetes_Cytophagaceae_Spirosoma
Bacteroidetes_Flavobacteriaceae_Chryseobacterium
Bacteroidetes_Sphingobacteriaceae_Mucilaginibacter
Bacteroidetes_Sphingobacteriaceae_Pedobacter
Firmicutes_Lactobacillaceae_Lactobacillus
Firmicutes_Paenibacillaceae_1_Paenibacillus
Proteobacteria_Caulobacteraceae_Brevundimonas
Proteobacteria_Enterobacteriaceae_Pantoea
Proteobacteria_Methylobacteriaceae_Methylobacterium
Proteobacteria_Moraxellaceae_Acinetobacter
Proteobacteria_Neisseriaceae_Snodgrassella
Proteobacteria_Orbaceae_Frischella
Proteobacteria_Oxalobacteraceae_Duganella
Proteobacteria_Oxalobacteraceae_Massilia
Proteobacteria_Pseudomonadaceae_Pseudomonas
Proteobacteria_Rhizobiaceae_Rhizobium
Proteobacteria_Sphingomonadaceae_Sphingomonas
Proteobacteria_Xanthomonadaceae_Luteibacter
Proteobacteria_Xanthomonadaceae_Stenotrophomonas
Proteobacteria_Xanthomonadaceae_Xanthomonas
Tenericutes_Entomoplasmataceae_Mesoplasma
-1
-0.8
-0.6
-0.4
-0.2
0
0.2
0.4
0.6
0.8
1

**Figure S2C**

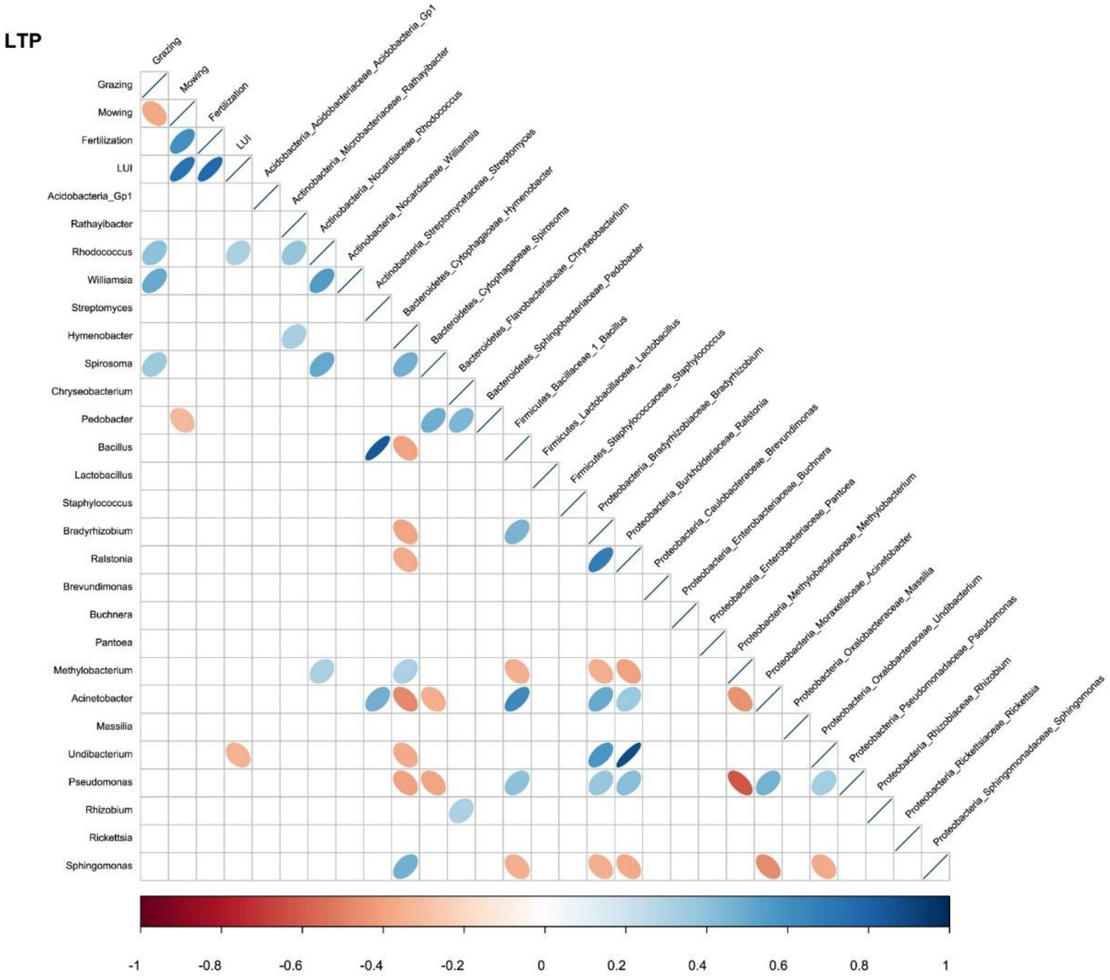

LTP
Grazing
Mowing
Fertilization
LUI
Acidobacteria_Acidobacteriaceae_Acidobacteria_Gp1
Actinobacteria_Microbacteriaceae_Rathayibacter
Actinobacteria_Nocardiaceae_Rhodococcus
Actinobacteria_Nocardiaceae_Williamsia
Actinobacteria_Streptomycetaceae_Streptomyces
Bacteroidetes_Cytophagaceae_Hymenobacter
Bacteroidetes_Cytophagaceae_Spirosoma
Bacteroidetes_Flavobacteriaceae_Chryseobacterium
Bacteroidetes_Sphingobacteriaceae_Pedobacter
Firmicutes_Bacillaceae_1_Bacillus
Firmicutes_Lactobacillaceae_Lactobacillus
Firmicutes_Staphylococcaceae_Staphylococcus
Proteobacteria_Bradyrhizobiaceae_Bradyrhizobium
Proteobacteria_Burkholderiaceae_Ralstonia
Proteobacteria_Caulobacteraceae_Brevundimonas
Proteobacteria_Enterobacteriaceae_Buchnera
Proteobacteria_Enterobacteriaceae_Pantoea
Proteobacteria_Methylobacteriaceae_Methylobacterium
Proteobacteria_Moraxellaceae_Acinetobacter
Proteobacteria_Oxalobacteraceae_Massilia
Proteobacteria_Oxalobacteraceae_Undibacterium
Proteobacteria_Pseudomonadaceae_Pseudomonas
Proteobacteria_Rhizobiaceae_Rhizobium
Proteobacteria_Rickettsiaceae_Rickettsia
Proteobacteria_Sphingomonadaceae_Sphingomonas
Grazing
Mowing
Fertilization
LUI
Acidobacteria_Gp1
Rathayibacter
Rhodococcus
Williamsia
Streptomyces
Hymenobacter
Spirosoma
Chryseobacterium
Pedobacter
Bacillus
Lactobacillus
Staphylococcus
Bradyrhizobium
Ralstonia
Brevundimonas
Buchnera
Pantoea
Methylobacterium
Acinetobacter
Massilia
Undibacterium
Pseudomonas
Rhizobium
Rickettsia
Sphingomonas
-1
-0.8
-0.6
-0.4
-0.2
0
0.2
0.4
0.6
0.8
1

**Figure S2D**

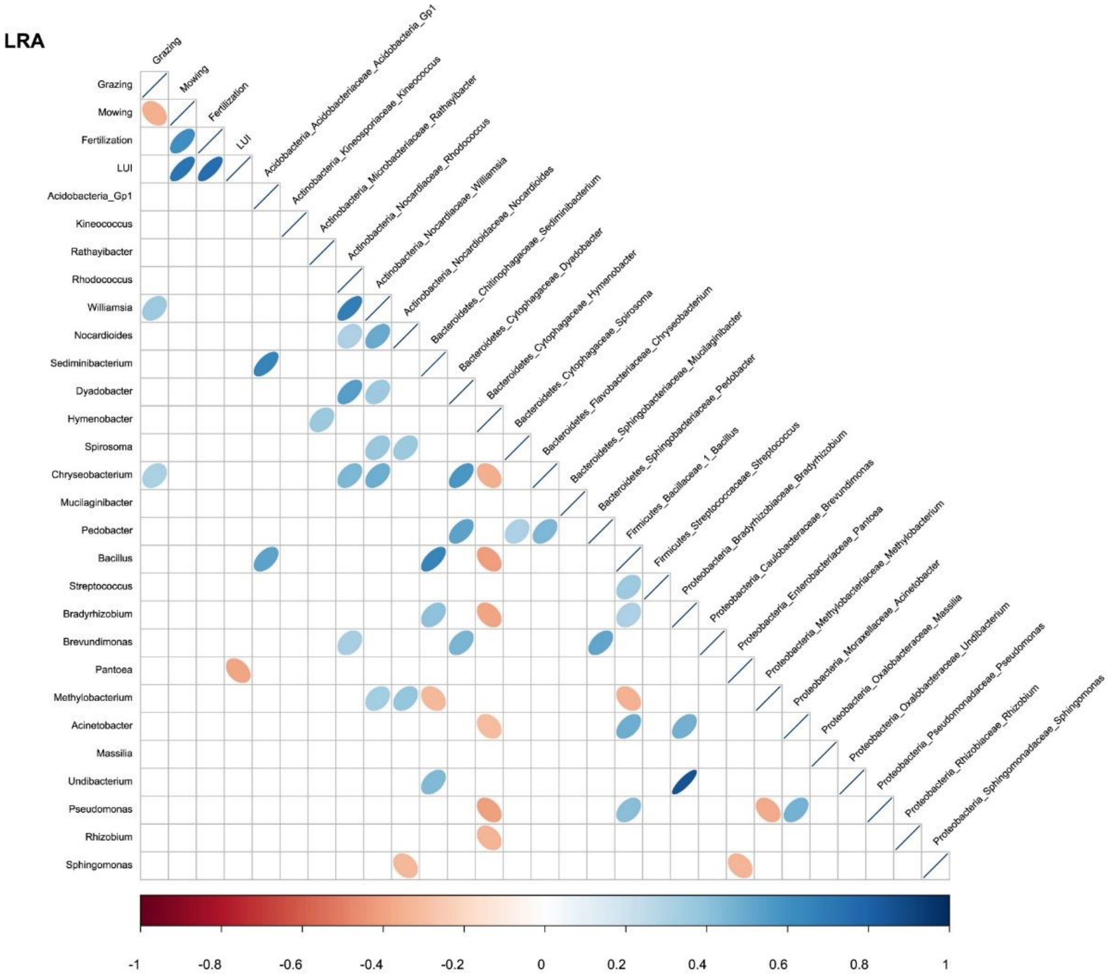
LRA
Grazing
Mowing
Fertilization
LUI
Acidobacteria_Acidobacteriaceae_Acidobacteria_Gp1
Actinobacteria_Kineosporiaceae_Kineococcus
Actinobacteria_Microbacteriaceae_Rathayibacter
Actinobacteria_Nocardiaceae_Rhodococcus
Actinobacteria_Nocardiaceae_Williamsia
Actinobacteria_Nocardioidaceae_Nocardioides
Bacteroidetes_Chitinophagaceae_Sediminibacterium
Bacteroidetes_Cytophagaceae_Dyadobacter
Bacteroidetes_Cytophagaceae_Hymenobacter
Bacteroidetes_Cytophagaceae_Spirosoma
Bacteroidetes_Flavobacteriaceae_Chryseobacterium
Bacteroidetes_Sphingobacteriaceae_Mucilaginibacter
Bacteroidetes_Sphingobacteriaceae_Pedobacter
Firmicutes_Bacillaceae_1_Bacillus
Firmicutes_Streptococcaceae_Streptococcus
Proteobacteria_Bradyrhizobiaceae_Bradyrhizobium
Proteobacteria_Caulobacteraceae_Brevundimonas
Proteobacteria_Enterobacteriaceae_Pantoea
Proteobacteria_Methylobacteriaceae_Methylobacterium
Proteobacteria_Moraxellaceae_Acinetobacter
Proteobacteria_Oxalobacteraceae_Massilia
Proteobacteria_Oxalobacteraceae_Undibacterium
Proteobacteria_Pseudomonadaceae_Pseudomonas
Proteobacteria_Rhizobiaceae_Rhizobium
Proteobacteria_Sphingomonadaceae_Sphingomonas
Grazing
Mowing
Fertilization
LUI
Acidobacteria_Gp1
Kineococcus
Rathayibacter
Rhodococcus
Williamsia
Nocardioides
Sediminibacterium
Dyadobacter
Hymenobacter
Spirosoma
Chryseobacterium
Mucilaginibacter
Pedobacter
Bacillus
Streptococcus
Bradyrhizobium
Brevundimonas
Pantoea
Methylobacterium
Acinetobacter
Massilia
Undibacterium
Pseudomonas
Rhizobium
Sphingomonas
-1
-0.8
-0.6
-0.4
-0.2
0
0.2
0.4
0.6
0.8
1

**Table S1**

| Phylum | Class | Order | Family | Genus | FRA(%) | FTP(%) | LRA(%) | LTP(%) |
|---|---|---|---|---|---|---|---|---|
| Proteobacteria | Gammaproteobacteria | Pseudomonadales | Pseudomonadaceae | Pseudomonas | 25.30 | 35.75 | 2.16 | 6.18 |
| Proteobacteria | Alphaproteobacteria | Sphingomonadales | Sphingomonadaceae | Sphingomonas | 6.45 | 8.18 | 23.87 | 19.38 |
| Proteobacteria | Gammaproteobacteria | Enterobacteriales | Enterobacteriaceae | Unclassified | 17.87 | 26.51 | 2.06 | 4.36 |
| Proteobacteria | Alphaproteobacteria | Rhizobiales | Methylobacteriaceae | Methylobacterium | 1.04 | 2.68 | 15.51 | 20.19 |
| Bacteroidetes | Cytophagia | Cytophagales | Cytophagaceae | Hymenobacter | 5.90 | 0.75 | 21.19 | 7.28 |
| Proteobacteria | Gammaproteobacteria | Unclassified | Unclassified | Unclassified | 1.32 | 0.55 | 0.11 | 12.56 |
| Proteobacteria | Alphaproteobacteria | Sphingomonadales | Sphingomonadaceae | Unclassified | 2.00 | 0.96 | 4.80 | 0.89 |
| Proteobacteria | Betaproteobacteria | Burkholderiales | Comamonadaceae | Unclassified | 1.22 | 1.40 | 4.32 | 1.19 |
| Firmicutes | Bacilli | Lactobacillales | Lactobacillaceae | Lactobacillus | 3.84 | 1.76 | 0.08 | 0.93 |
| Proteobacteria | Gammaproteobacteria | Pseudomonadales | Moraxellaceae | Acinetobacter | 5.24 | 0.34 | 0.24 | 0.57 |
| Bacteroidetes | Cytophagia | Cytophagales | Cytophagaceae | Spirosoma | 0.49 | 0.24 | 3.87 | 1.68 |
| Proteobacteria | Betaproteobacteria | Burkholderiales | Oxalobacteraceae | Duganella | 1.47 | 3.65 | 0.09 | 0.26 |
| Bacteroidetes | Sphingobacteriia | Sphingobacteriales | Sphingobacteriaceae | Pedobacter | 0.96 | 1.66 | 1.32 | 0.67 |
| Actinobacteria | Actinobacteria | Actinomycetales | Microbacteriaceae | Unclassified | 0.48 | 0.53 | 1.84 | 1.45 |
| Proteobacteria | Alphaproteobacteria | Rhizobiales | Unclassified | Unclassified | 0.58 | 0.09 | 3.09 | 0.28 |
| Bacteroidetes | Flavobacteriia | Flavobacteriales | Flavobacteriaceae | Chryseobacterium | 0.48 | 2.43 | 0.38 | 0.53 |
| Proteobacteria | Alphaproteobacteria | Rhizobiales | Rhizobiaceae | Rhizobium | 0.64 | 1.02 | 1.38 | 0.73 |
| Proteobacteria | Gammaproteobacteria | Pseudomonadales | Moraxellaceae | Unclassified | 2.97 | 0.01 | 0.04 | 0.22 |
| Tenericutes | Mollicutes | Entomoplasmatales | Unclassified | Unclassified | 3.09 | 0.05 | 0.02 | 0.02 |
| Tenericutes | Mollicutes | Entomoplasmatales | Entomoplasmataceae | Mesoplasma | 2.94 | 0.05 | 0.02 | 0.03 |
| Proteobacteria | Alphaproteobacteria | Rickettsiales | Rickettsiaceae | Rickettsia | 0.85 | 0.02 | 0.08 | 1.88 |
| Proteobacteria | Betaproteobacteria | Burkholderiales | Oxalobacteraceae | Massilia | 0.96 | 0.63 | 0.60 | 0.49 |
| Actinobacteria | Actinobacteria | Actinomycetales | Nocardiaceae | Rhodococcus | 0.07 | 0.18 | 0.68 | 1.54 |
| Proteobacteria | Unclassified | Unclassified | Unclassified | Unclassified | 0.36 | 0.16 | 0.53 | 1.09 |
| Proteobacteria | Alphaproteobacteria | Rhodospirillales | Acetobacteraceae | Unclassified | 0.18 | 1.26 | 0.41 | 0.18 |
| Proteobacteria | Gammaproteobacteria | Xanthomonadales | Xanthomonadaceae | Stenotrophomonas | 0.03 | 1.70 | 0.06 | 0.16 |
| Proteobacteria | Gammaproteobacteria | Enterobacteriales | Enterobacteriaceae | Arsenophonus | 1.82 | 0.01 | 0.03 | 0.09 |
| Proteobacteria | Alphaproteobacteria | Rhizobiales | Rhizobiaceae | Unclassified | 0.34 | 1.06 | 0.20 | 0.18 |
| Firmicutes | Bacilli | Bacillales | Staphylococcaceae | Staphylococcus | 0.65 | 0.03 | 0.07 | 0.87 |
| Firmicutes | Bacilli | Lactobacillales | Streptococcaceae | Lactococcus | 1.40 | 0.01 | 0.05 | 0.03 |
| Proteobacteria | Alphaproteobacteria | Rhizobiales | Aurantimonadaceae | Unclassified | 0.13 | 0.19 | 0.67 | 0.36 |
| Proteobacteria | Gammaproteobacteria | Enterobacteriales | Enterobacteriaceae | Buchnera | 0.01 | 0.03 | 0.02 | 1.09 |
| Firmicutes | Bacilli | Lactobacillales | Leuconostocaceae | Leuconostoc | 0.88 | 0.01 | 0.05 | 0.11 |
| Proteobacteria | Alphaproteobacteria | Caulobacterales | Caulobacteraceae | Brevundimonas | 0.08 | 0.28 | 0.34 | 0.34 |
| Actinobacteria | Actinobacteria | Actinomycetales | Microbacteriaceae | Rathayibacter | 0.04 | 0.07 | 0.39 | 0.53 |
| Firmicutes | Bacilli | Bacillales | Paenibacillaceae | Paenibacillus | 0.47 | 0.44 | 0.01 | 0.03 |
| Bacteroidetes | Cytophagia | Cytophagales | Cytophagaceae | Dyadobacter | 0.15 | 0.35 | 0.33 | 0.11 |
| Proteobacteria | Betaproteobacteria | Burkholderiales | Alcaligenaceae | Unclassified | 0.03 | 0.61 | 0.20 | 0.08 |
| Proteobacteria | Betaproteobacteria | Burkholderiales | Unclassified | Unclassified | 0.06 | 0.53 | 0.15 | 0.17 |
| Proteobacteria | Gammaproteobacteria | Legionellales | Coxiellaceae | Diplorickettsia | 0.75 | 0.00 | 0.05 | 0.01 |
| Proteobacteria | Gammaproteobacteria | Xanthomonadales | Xanthomonadaceae | Xanthomonas | 0.17 | 0.55 | 0.03 | 0.04 |
| Firmicutes | Bacilli | Bacillales | Bacillaceae | Bacillus | 0.04 | 0.03 | 0.13 | 0.52 |
| Actinobacteria | Actinobacteria | Actinomycetales | Kineosporiaceae | Unclassified | 0.04 | 0.01 | 0.49 | 0.16 |
| Bacteroidetes | Sphingobacteriia | Sphingobacteriales | Sphingobacteriaceae | Mucilaginibacter | 0.09 | 0.15 | 0.28 | 0.15 |
| Actinobacteria | Actinobacteria | Actinomycetales | Nocardioidaceae | Nocardioides | 0.02 | 0.01 | 0.35 | 0.28 |
| Bacteroidetes | Sphingobacteriia | Sphingobacteriales | Chitinophagaceae | Unclassified | 0.08 | 0.02 | 0.34 | 0.19 |
| Proteobacteria | Betaproteobacteria | Burkholderiales | Oxalobacteraceae | Undibacterium | 0.04 | 0.02 | 0.13 | 0.40 |
| Actinobacteria | Actinobacteria | Actinomycetales | Geodermatophilaceae | Geodermatophilus | 0.02 | 0.02 | 0.26 | 0.29 |
| Actinobacteria | Actinobacteria | Actinomycetales | Nocardiaceae | Williamsia | 0.02 | 0.02 | 0.25 | 0.30 |
| Proteobacteria | Alphaproteobacteria | Rhizobiales | Bradyrhizobiaceae | Bradyrhizobium | 0.04 | 0.03 | 0.12 | 0.36 |
| Proteobacteria | Gammaproteobacteria | Oceanospirillales | Halomonadaceae | Unclassified | 0.38 | 0.00 | 0.01 | 0.11 |
| Actinobacteria | Actinobacteria | Actinomycetales | Unclassified | Unclassified | 0.03 | 0.01 | 0.21 | 0.23 |
| Bacteroidetes | Sphingobacteriia | Sphingobacteriales | Chitinophagaceae | Sediminibacterium | 0.06 | 0.03 | 0.16 | 0.23 |
| Proteobacteria | Alphaproteobacteria | Rhizobiales | Aurantimonadaceae | Aureimonas | 0.04 | 0.04 | 0.19 | 0.18 |
| Acidobacteria | Acidobacteriia | Acidobacteriales | Acidobacteriaceae | Acidobacteria_Gp1 | 0.07 | 0.01 | 0.09 | 0.25 |
| Actinobacteria | Actinobacteria | Actinomycetales | Kineosporiaceae | Kineococcus | 0.03 | 0.04 | 0.21 | 0.14 |
| Proteobacteria | Betaproteobacteria | Neisseriales | Neisseriaceae | Snodgrassella | 0.02 | 0.21 | 0.16 | 0.02 |
| Bacteroidetes | Flavobacteriia | Flavobacteriales | Flavobacteriaceae | Flavobacterium | 0.13 | 0.04 | 0.17 | 0.06 |

**Table S2**

| ASV-ID | Phylum | Class | Order | Family | Genus | FRA( %) | FTP (%) | LRA (%) | LTP (%) |
|---|---|---|---|---|---|---|---|---|---|
| ASV4 | Proteobacteria | Gammaproteobacteria | Pseudomonadales | Pseudomonadaceae | *Pseudomonas* | 10.22 | 20.24 | 2.18 | 4.14 |
| ASV5 | Proteobacteria | Alphaproteobacteria | Sphingomonadales | Sphingomonadaceae | *Sphingomonas* | 3.93 | 4.64 | 14.60 | 12.42 |
| ASV3 | Proteobacteria | Gammaproteobacteria | Enterobacteriales | Enterobacteriaceae | *Pantoea* | 5.32 | 22.79 | 2.31 | 3.48 |
| ASV7 | Proteobacteria | Gammaproteobacteria | Pseudomonadales | Pseudomonadaceae | *Pseudomonas* | 9.25 | 4.52 | 0.40 | 1.69 |
| ASV16 | Proteobacteria | Alphaproteobacteria | Sphingomonadales | Sphingomonadaceae | *Sphingomonas* | 2.32 | 0.87 | 4.25 | 0.59 |
| ASV12 | Proteobacteria | Alphaproteobacteria | Rhizobiales | Methylobacteriaceae | *Methylobacterium* | 0.28 | 0.75 | 2.32 | 3.96 |
| ASV58 | Proteobacteria | Alphaproteobacteria | Sphingomonadales | Sphingomonadaceae | *Sphingomonas* | 0.25 | 0.28 | 0.84 | 0.85 |
| ASV69 | Proteobacteria | Alphaproteobacteria | Sphingomonadales | Sphingomonadaceae | *Sphingomonas* | 0.19 | 0.66 | 0.56 | 0.64 |

**Table S3**

| Plant Tissue | Domain | Phylum | Class | p-value | ALB (%) | HAI (%) | SCH (%) |
|---|---|---|---|---|---|---|---|
| Flowers *R. acris* | Bacteria | Bacteroidetes | Cytophagia | 0.040* | 7.86 | 2.35 | 2.47 |
| Flowers *R. acris* | Bacteria | Bacteroidetes | Flavobacteriia | 0.025* | 0.41 | 0 | 0.87 |
| Flowers *R. acris* | Bacteria | Proteobacteria | Unclassified | < 0.001*** | 0 | 0.57 | 0 |
| Flowers *T. pratense* | Bacteria | Bacteroidetes | Cytophagia | 0.032* | 0.49 | 0.11 | 2.15 |
| Leaves *R. acris* | Bacteria | Firmicutes | Bacilli | < 0.001*** | 0 | 0 | 0.23 |
| Leaves *T. pratense* | Bacteria | Firmicutes | Bacilli | 0.002** | 0.71 | 5.87 | 1.31 |
| Leaves *T. pratense* | Bacteria | Bacteroidetes | Cytophagia | 0.015* | 10.93 | 2.59 | 7.34 |

**Table S4**

| Plant Tissue | Distance Metrics | $r^2$ | p-value |
|---|---|---|---|
| Flowers *R. acris* | Unweighted UniFrac | 0.01 | 0.008** |
| | Weighted UniFrac | 0.01 | 0.203 |
| | Bray-Curtis Dissimilarity | 0.01 | 0.222 |
| Flowers *T. pratense* | Unweighted UniFrac | 0.02 | 0.076 |
| | Weighted UniFrac | 0.01 | 0.482 |
| | Bray-Curtis Dissimilarity | 0.02 | 0.094 |
| Leaves *R. acris* | Unweighted UniFrac | 0.01 | 0.012* |
| | Weighted UniFrac | 0.01 | 0.262 |
| | Bray-Curtis Dissimilarity | 0.01 | 0.081 |
| Leaves *T. pratense* | Unweighted UniFrac | 0.02 | 0.015* |
| | Weighted UniFrac | 0.02 | 0.128 |
| | Bray-Curtis Dissimilarity | 0.02 | 0.082 |

### Table S5A

| Domain | Phylum | Class | Order | Family | p-value | low LUI (%) | high LUI (%) |
|---|---|---|---|---|---|---|---|
| Bacteria | Verrucomicrobia | Opitutae | Opitutales | Opitutaceae | 0.016 | **0.01255** | 0.00193 |
| Bacteria | Actinobacteria | Actinobacteria | Acidimicrobiales | Iamiaceae | 0.023 | **0.00364** | 0.00074 |
| Bacteria | Proteobacteria | Deltaproteobacteria | Desulfuromonadales | Geobacteraceae | 0.024 | **0.00344** | 0.00015 |
| Bacteria | Actinobacteria | Actinobacteria | Actinomycetales | Thermomonosporaceae | 0.029 | **0.02838** | 0.00423 |
| Bacteria | Proteobacteria | Deltaproteobacteria | Myxococcales | Unclassified | 0.030 | **0.03586** | 0.00777 |
| Bacteria | Chloroflexi | Caldilineae | Caldilineales | Caldilineaceae | 0.040 | **0.00763** | 0.00099 |
| Bacteria | Proteobacteria | Deltaproteobacteria | Myxococcales | Myxococcaceae | 0.043 | **0.00708** | 0.00004 |
| Bacteria | Proteobacteria | Gammaproteobacteria | Oceanospirillales | Halomonadaceae | 0.045 | **0.59303** | 0.09853 |
| Bacteria | Tenericutes | Mollicutes | Entomoplasmatales | Entomoplasmataceae | 0.047 | 0.79129 | **6.15646** |
| Bacteria | Proteobacteria | Alphaproteobacteria | Rhodospirillales | Acetobacteraceae | 0.048 | **1.45008** | 0.57852 |
| Bacteria | Verrucomicrobia | Subdivision3 | Unclassified | Unclassified | 0.049 | **0.11861** | 0.02429 |

### Table S5B

| Domain | Phylum | Class | Order | Family | p-value | low LUI (%) | high LUI (%) |
|---|---|---|---|---|---|---|---|
| Bacteria | Bacteroidetes | Cytophagia | Cytophagales | Cytophagaceae | 0.012 | 0.84717 | **1.90630** |
| Bacteria | Proteobacteria | Alphaproteobacteria | Caulobacterales | Caulobacteraceae | 0.014 | 0.11553 | **0.43665** |
| Bacteria | Actinobacteria | Actinobacteria | Actinomycetales | Nocardioidaceae | 0.022 | 0.01405 | **0.04425** |
| Bacteria | Bacteroidetes | Sphingobacteriia | Sphingobacteriales | Rhodothermaceae | 0.023 | 0.00000 | **0.00129** |
| Bacteria | Actinobacteria | Actinobacteria | Actinomycetales | Unclassified | 0.027 | 0.00608 | **0.01900** |
| Bacteria | Proteobacteria | Gammaproteobacteria | Xanthomonadales | Xanthomonadaceae | 0.039 | **2.84370** | 1.39912 |
| Bacteria | Actinobacteria | Actinobacteria | Solirubrobacterales | Solirubrobacteraceae | 0.045 | 0.00023 | **0.00140** |
| Bacteria | Proteobacteria | Betaproteobacteria | Burkholderiales | Alcaligenaceae | 0.048 | **0.67329** | 0.40087 |

### Table S5C

| Domain | Phylum | Class | Order | Family | p-value | low LUI (%) | high LUI (%) |
|---|---|---|---|---|---|---|---|
| Bacteria | Proteobacteria | Gammaproteobacteria | Enterobacteriales | Unclassified | 0.023 | **0.00406** | 0.00078 |
| Bacteria | Proteobacteria | Gammaproteobacteria | Enterobacteriales | Enterobacteriaceae | 0.026 | **5.82054** | 1.23627 |
| Bacteria | Proteobacteria | Alphaproteobacteria | Sphingomonadales | Sphingomonadaceae | 0.039 | 24.34726 | **30.16829** |
| Bacteria | Proteobacteria | Alphaproteobacteria | Rhizobiales | Rhizobiaceae | 0.044 | **3.23699** | 1.30514 |
| Bacteria | Actinobacteria | Actinobacteria | Actinomycetales | Nocardiopsaceae | 0.047 | **0.00178** | 0.00000 |

### Table S5D

| Domain | Phylum | Class | Order | Family | p-value | low LUI (%) | high LUI (%) |
|---|---|---|---|---|---|---|---|
| Bacteria | Proteobacteria | Alphaproteobacteria | Rhodospirillales | Acetobacteraceae | 0.003 | **0.72818** | 0.35348 |
| Bacteria | Actinobacteria | Actinobacteria | Actinomycetales | Nocardiaceae | 0.009 | 1.00057 | **2.46234** |
| Bacteria | Proteobacteria | Alphaproteobacteria | Rhodobacterales | Unclassified | 0.023 | **0.00141** | 0.00000 |
| Bacteria | Bacteroidetes | Flavobacteriia | Flavobacteriales | Cryomorphaceae | 0.024 | **0.00530** | 0.00036 |
| Bacteria | Firmicutes | Bacilli | Lactobacillales | Aerococcaceae | 0.024 | **0.04852** | 0.00678 |
| Bacteria | Actinobacteria | Actinobacteria | Gaiellales | Gaiellaceae | 0.035 | **0.07515** | 0.02493 |
| Bacteria | Actinobacteria | Actinobacteria | Actinomycetales | Sanguibacteraceae | 0.035 | **0.02366** | 0.00008 |
| Bacteria | Armatimonadetes | Armatimonadia | Armatimonadales | Armatimonadaceae | 0.040 | **0.03579** | 0.01319 |
| Bacteria | Firmicutes | Clostridia | Clostridiales | Lachnospiraceae | 0.041 | **0.03351** | 0.00979 |
| Bacteria | Actinobacteria | Actinobacteria | Rubrobacterales | Rubrobacteraceae | 0.049 | **0.00613** | 0.00055 |
| Bacteria | Proteobacteria | Deltaproteobacteria | Myxococcales | Cystobacteraceae | 0.050 | **0.01044** | 0.00113 |
